# Probing other’s Presence: Probabilistic Inference Across Brain Scales Reveals Enhanced Excitatory Synaptic Efficacy

**DOI:** 10.1101/2024.09.09.612006

**Authors:** Amirhossein Esmaeili, Marie Demolliens, Marjolein Viersen, Abolfazl Ziaeemehr, Faical Isbaine, Pascal Huguet, Frank Zaal, Viktor Jirsa, Driss Boussaoud, Meysam Hashemi

**Affiliations:** Aix Marseille University, INSERM, INS, Inst Neurosci Syst, Marseille, France; Laboratory of Social and Cognitive Psychology, Clermont-Ferrand University, France; Department of Human Movement Sciences, University Medical Center Groningen, Groningen, Netherlands; Department of Neurosurgery, Emory University, Atlanta, United States

**Keywords:** Social facilitation and impairment effects, Attentional modulation, Multi-scale inference, Deep neural density estimators, Effective connectivity, Synaptic efficacy

## Abstract

The presence of conspecifics is a fundamental and arguably invariant prerequisite of social cognition across numerous animal species. While the influence of social presence on behavior has been among the focal points of investigation in social psychology for over a century, its underlying neural mechanisms remain largely unexplored. Here, we attempt to bridge this gap by investigating how the presence of conspecifics changes synaptic efficacy from measurements across spatiotemporal brain scales, and how such changes could lead to modulations of task performance in monkeys and humans. In monkeys performing an association learning task, social presence increased excitatory synaptic efficacy in attention-oriented regions dorsolateral prefrontal cortex and anterior cingulate cortex. In humans performing a visuomotor task, the presence of conspecifics facilitated performance in one of the subject groups, and this facilitation was linked to enhanced excitatory synaptic efficacy within the dorsal and ventral attention networks. We propose that presence-induced improvements in task performance arise from attentional modulation mediated by changes in excitatory synaptic efficacy across three spatiotemporal brain scales, namely, micro-scale (single neurons), meso-scale (cortical columns) and macro-scale (whole-brain). Our findings from Bayesian learning converge to establish a novel, multi-scale framework for understanding the neural underpinnings of social presence effects. This probabilistic framework offers a fresh perspective on social presence research, and lays the groundwork for future investigations into the complex interplay between social presence, neural dynamics, and behavior.

## 1. Introduction

Social isolation wreaks havoc upon the brain [1]. Social cognition –with cognition going beyond the scope of simple perception–, as with non-social cognition, hinges first and foremost on the capacity to perceive relevant stimuli in the environment, namely social agents. Perception of members of the same species, formally referred to as conspecifics, is therefore an integral –and seemingly primordial– prerequisite for many forms of interaction, from cooperation to competition [2]. Despite the tremendous effort for delineating the neural correlates of social cognition [3, 4], the literature surrounding the exact neurobiological mechanism of the perception/awareness of others’ mere presence –the most fundamental invariant aspect of social cognition in many, if not all, animal species– is surprisingly scant. This sparsity might potentially stem from the difficulty of decomposition of presence ‘stimuli’. While social perception is undoubtedly a pivotal part of cognition in many mammalian species, it would stand to reason that such a perception in each species would be predicated upon the sensory modality towards which they are most adapted. For instance, humanoid primates rely primarily on vision, while rodents rely on olfaction for recognition of other social agents [5]. Decomposition of neural correlates of simple ‘presence’ from other aspects of perception might prove to be an intricate and highly complex endeavor. The closest –and potentially feasible– logical avenue of investigation would be to analyze the behavioral modulations resulting from the mere presence of conspecifics. Fortuitously, this avenue of research is among the oldest and most well-explored areas of experimental social psychology [6, 7, 8, 9]. Indeed, social presence has been repeatedly shown to modulate behavior in a variety of animal species including human and non-human primates [10, 11, 12, 13], and could be conceived as the closest avenue of inquiry for investigating mere presence effects. Although such a modulation is commonly referred to as ‘social facilitation’, mere presence may actually either improve (social facilitation) or impair (social impairment) performance, compared to isolation depending on task complexity [8] and the more or less threatening and/or evaluative nature of social presence [7]. While social facilitation/impairment (SFI) effects have been relatively well-researched from a behavioral perspective, much remains to be understood regarding its neural underpinnings [10].

Two main frameworks have been proposed regarding the underlying mechanisms of SFI effects. Zajonc (1965) was the first to notice that the presence of others (as observers or coactors) typically facilitates performance on easy or well-learned tasks and impairs performance on difficult or poorly learned tasks. On the basis of the Hull-Spence behaviorist theory of learning, conditioning, and motivation (well accepted in the 1950s and 1960s), Zajonc suggested that the mere presence of conspecifics energizes the emission of the dominant (habitual/prepotent or automatic) responses. According to the Hull-Spence theory, the energization of dominant responses indeed improves performance in well-learned tasks in which, by definition, correct responses are dominant and deteriorates performance in poorly learned tasks in which errors are the most likely responses. Zajonc’s view of SFI effects found support in many studies using very different species whose dominant responses–whether correct or incorrect–increased under social presence, compared with isolation. However, although Zajonc’s classic view remains the most common interpretation of SFI effects (see also [14] for a motivational account close to Zajonc’s), there is a large body of evidence that these effects can also involve attentional mechanisms [6], at least in humans and nonhuman primates, either by facilitating attentional focusing or by undermining cognitive control (which heavily relies on executive attention) in self-threatening circumstances [13]. As for the neural correlates of other’s presence, the scant number of existing studies provide further support for modulation of attentional regions/networks [15, 16, 10, 11]. However, the specific neural mechanisms underlying SFI effects remain unclear. Given that some recent inquiries have pointed to the existence of context-sensitive neural populations in prefrontal regions that are preferentially active either during conspecific’s presence or absence, it would stand to reason that the representative characteristics of the recruited neural ensembles during others’ presence would differ from those observed during isolation [10]. Upon initial glance, empirical investigations of these characteristics in higher primates face seemingly insurmountable challenges. For one, the timescale of social facilitation effects –by definition– precludes the possibility of in-vitro analyses in non-human primates. Second, neuroimaging techniques in humans only provide derived measures of neural activity (such as the scalp potential generated by the collective neuronal activity of large populations in EEG or hemodynamic activity in fMRI), making it challenging to directly reveal potential hidden causal mechanisms at the synaptic level. Through the choice of appropriate computational models and probabilistic inference, however, one could reasonably approximate the relation between measured neural activities and specific model configurations (i.e., generative parameters). Dynamic causal modeling (DCM; [17]) is a well-established statistical framework to reveal the hidden causal mechanisms within neural subsystems –at lower levels– (e.g. synapses), from measurements such as event-related potentials (ERPs) or EEG. In other words, such Bayesian approach allows for creation and testing of in-silico hypotheses regarding the causal mechanisms within biological neural networks and their contribution to various cognitive processes or behaviors. In this context, the causal influence exerted by one neural subsystem on another is commonly referred to as effective connectivity [18, 19]. A large body of work supports the associations between attentional modulation in the brain and changes in effective connectivity [20]. For instance, evidence points to associations between motion-oriented area V5 and posterior parietal cortex in attending to actions [20], prefrontal and premotor cortices during attention to visual motion [21], and between visual cortex and medial temporal lobe [22]. There has even been evidence supporting a positive relation between increased attentional demands of a task and enhanced effective connectivity within the descending attention pathway [23]. This attentional modulation is supposedly achieved via optimization of neural communication at the synaptic level through selective modification of synaptic efficacy (SE), which serves as the neurobiological proxy for modulations in effective connectivity [24]. It is important to note that synaptic inputs can stem from either excitatory or inhibitory presynaptic cells, modulations of either of which could bear strong consequences for Excitation/Inhibition (E/I) balance (i.e. in a degenerate manner [25]). The E/I balance in neural networks is critical for maintaining stable network activity, enabling efficient information processing. Moreover, disruptions in E/I balance are commonly linked to cognitive deficits and various neurological disorders, including ASD [26].

We hypothesize that modulation of brain dynamics –and their subsequent influence on behavior– due to mere presence could be reliably detected as variations of SE across principal spatiotemporal scales of the primate brain. To test this hypothesis, we employed the framework of DCM based on the data from our facilitation tasks involving non-human and human primates (which are done either in the presence or absence of a social conspecific), in order to link the modulations of task performance to adaptations of SE across three brain scales, namely microscale (single neurons), mesoscale (cortical columns), and macroscale (whole-brain).

Although challenging, conducting DCM across brain scales can provide a more comprehensive understanding of neural dynamics, potentially revealing a universal principle of brain function. When leveraged with advanced probabilistic machine learning techniques [27, 28, 29], this approach can operate across brain scales, enhancing the accuracy of models inferring the neural mechanisms underlying social facilitation, as we demonstrate in this study. We achieved this by estimating synaptic efficacies across said scales using state-of-the-art deep neural density estimators [30, 31, 29]. Tailored to Bayesian learning, a family of these generative models called normalizing flows [31] applies a series of invertible mappings in order to efficiently transform a simple base distribution into any complex target distribution. We employ this framework in the current study in order to calculate the distribution of synaptic efficacies, given the summary statistics of empirical evidence. This is motivated by the theory of structured flows on manifold ([32, 33]), which posits that brain dynamics and behavior are both constrained to low-dimensional subspaces and are topologically equivalent.

In the following we will outline the association-learning and lateral-interception tasks conducted in monkeys and humans, respectively. Subsequently, we will demonstrate how presence modulates behavioral dynamics and identify neural correlates of this modulation in the previously mentioned tasks. Finally, we use these neural correlates as informative data features, in order to agnostically estimate synaptic efficacies across three spatiotemporal brain scales, namely microscale (single neurons), mesoscale (cortical columns), and macroscale (whole-brain).

## 2. Results

### 2.1. Association learning in presence vs absence

Monkeys were trained on a touchscreen task to associate abstract cues with specific targets. The experiment consisted of two conditions: social presence and isolation (details on housing and experimental setup provided in Supplementary subsection 6.1). During social presence trials (Figure 1**A**), monkeys faced each other, with each taking on the role of either the actor (performing the task) or the passive spectator (observing the actor). Only the actor interacted with the touchscreen, receiving rewards for correct choices. Spectators remained unrewarded and never participated in the same session as actors to prevent observational learning. Conversely, during isolation trials (Figure 1**B**), monkeys performed the task alone, deprived of any visual, auditory, or olfactory contact with their conspecific. The behavioral data obtain through this task indicated a social facilitation effect in both monkeys (see Figure S2, generated from [10]).

**Figure 1.**
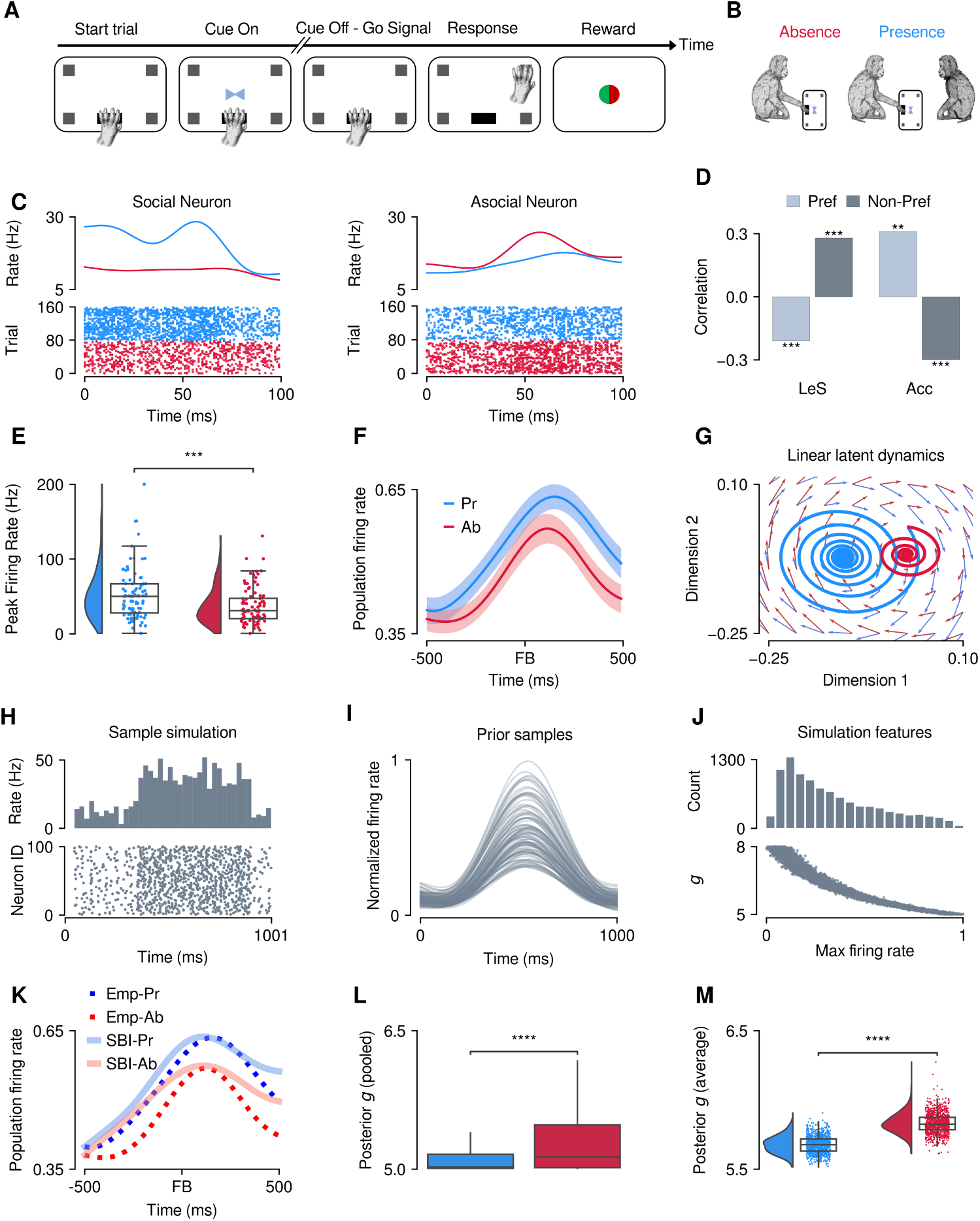
Inference of microscale synaptic efficacy. **A**) Association learning task outline in Monkeys, during which subjects had to associate which of the four corners of the touchpad was associated with the cue in order to obtain a food reward. **B**) The two experimental conditions in the association learning task. **C**) Sample firing rates (top) and spike-rasters (bottom) of social (left) and asocial (right) neurons. **D**) Correlation of the firing rate of social/asocial neurons with task performance. **E**) Peak firing rate of all categorized neurons in the presence condition (blue) compared to the absence (red) (*p <* 0.001). **F**) Average firing rate of the same neurons as in panel E. **G**) Linear latent dynamics of single-neuron firing rates in the two experimental conditions. **H**) Sample firing rate histogram (top) and spike-raster of simulations of the balanced spiking network. **I**) Sample firing rate of simulations with random values of parameter *g* from prior distribution. **J**) Peak firing rate of the model based on varying values of SE (bottom), and the histogram of said peaks (top). This peak firing rate is used for training the deep neural density estimators. **K**) Empirical (dotted lines) and posterior predictive fits (solid lines) of average firing rates in presence/absence. **L**) Pooled distribution of inferred SE across all neurons (*p <* 0.0001). **M**) Distribution of inferred SE from the average firing rates of the recorded regions (*p <* 0.0001).

### 2.2. Presence/Absence oriented neural ensembles

We discovered neural subpopulations in both dorsolateral prefrontal cortex (dlPFC) and anterior cingulate cortex (ACC) that were preferentially active under one of the two experimental conditions [10]. Neurons that fired more during the presence condition were thus categorized as ‘social neurons’, and those that fired more under social isolation, as ‘asocial neurons’. This firing rate strongly correlated with both learning speed and accuracy (trial-to-criterion) in the neurons’ preferred conditions (social neurons under presence; asocial neurons under isolation), while exhibiting negative correlations in the non-preferred conditions (Learning speed (Les): *p <* 0.001 for both conditions, Accuracy (Acc): *p <* 0.01 in preferred, and *p <* 0.001 in non-preferred conditions; Figure 1**C**, **D**). In addition, we found that the peak firing rates of neurons in both of these subpopulations were significantly elevated in the presence condition compared to the absence condition (*p <* 0.001; Figure 1**E**, **F**). To validate these findings, we fitted the firing rate of all neurons in each condition to a linear dynamical system to infer the latent dynamics (Methods subsection 5.13) in the presence and absence conditions. By sampling from these learned dynamics, we observed that changes in firing rate could be seen as variations in the decay rate (relaxation of population activity; [34, 35]), which reflects the rate of convergence to the stable fixed point of the linear system (Figure 1**G**). Functionally speaking, this framework enabled us to estimate the rate by which the linear system (i.e. neural activity) returns to its equilibrium after a perturbation in its measured output, revealing a slower decay rate in the presence condition, compared to absence, which could hint at a potential neuromodulatory role for others’ presence. However, none of the above findings can directly elucidate how SE is altered in the presence of others.

### 2.3. Enhanced effective connectivity during conspecific presence at microscale

Understanding the underlying neurobiological mechanisms that drive changes in neural firing rates is crucial for unraveling the complexities of neural computation and communication. Task-oriented single neurons exhibit specific firing rate patterns that are thought to influence the coarse-grain network dynamics. However, delineating the precise relationship between individual neuron activity and network dynamics remains non-trivial. By using a balanced spiking neural network model [36], we aim to bridge this gap. This model allows us to simulate and analyze how changes in SE within the network can lead to changes in firing rates. The balanced nature of the model ensures that excitatory and inhibitory inputs are finely tuned, mirroring the conditions found in biological neural networks. This model (see Equation 1) comprises of one excitatory (*n* = 10000) and one inhibitory (*n* = 2500) subpopulation of neurons (Figure 1**H**). The strength of connections between the two subpopulations is scaled by negative *g*, where *g* parameter denotes the ratio of inhibitory to excitatory weights across the entire population. In other words, higher values of *g* in this model translate to lower overall (regional) network activity (Figure 1**J**). To estimate the posterior distribution of parameter *g*, we trained deep neural density estimators (called neural spline flows; [31], Methods subsection 5.14) on the maximum firing rate of the excitatory subpopulation from *n* = 10000 random simulations (Figure 1**I**, **J**). After training, empirical data were used to estimate connectivity values that best represent the averaged firing rate in each experimental condition. By random sampling from the estimated posterior distribution, we achieved a close fit around the peaks while maintaining a consistent baseline (Figure 1**K**). We found significant decreases in parameter *g* not only in condition averages, but also when comparing conditions across all neurons (*p <* 0.0001 for both pooled, and average firing rates; Figure 1**L**, **M**). In sum, these findings demonstrate that conspecific’s presence leads to an increase in effective connectivity between single neurons in attention-oriented regions.

### 2.4. Increased local recurrent excitation during conspecific presence at mesoscale

A global change to effective connectivity between single neurons does not inform us of the intricacies of how connectivity between different cortical subpopulations might be altered within a cortical column. To this end, we computed ERPs during presence and absence conditions. We observed that not only the ERPs in the presence condition possess a significantly higher peak compared to the absence condition (*p <* 0.001; Figure 2**A**, **B**), but also that the ratio of ERP peaks between conditions was correlated with the ratio of behavioral performance (accuracy per session) between conditions (*ρ_pearson_* = 0.33; Figure 2**C**). This finding is further validated by the clear separation of the embeddings of ERPs (Methods subsection 5.5) between the experimental conditions (Figure 2**D**). Evoked potentials typically measure synaptic activity at the population level, making it computationally implausible to infer the synaptic connection between every possible cortical subpopulation. Our model of choice is a modified derivation of the Jansen-Rit neural mass model [37, 38] –capable of generating evoked potentials (Figure 2**E**)– with three subpopulations comprising pyramidal neurons (PNs), interneurons (INs), and stellate cells (SCs), which provides a balance between computational efficiency and biological plausibility. The subpopulations are connected by four effective connectivity parameters *g*_1_*_−_*_4_. The biological plausibility of this model, however, translates to model degeneracy, where different sets of input parameters could lead to identical model output. By fixing the model time-scales at biological values (i.e., placing informative prior in Bayesian setting), we therefore trained neural density estimators on peak evoked values obtained from *n* = 10000 random simulations to approximate the joint posterior distribution of effective connectivities. Our inferred distribution of peak evoked activity –in each condition– shares similar characteristics with that of the empirical data, up to the second order of statistical moments (*p <* 0.001; Figure 2**F**). To further validate the estimation, we generated nonlinear latent dynamics from the learned connectivity values for each experimental session/condition (Figure 2**G**). These dynamics collectively formed the calculated embedding manifold, illustrating higher evoked activity in the presence condition compared to absence (Figure 2**H**). Pooling the connectivity distributions across all sessions, we found a significant difference between the experimental conditions only with respect to the parameter *g*_2_, which provides local recurrent excitation to pyramidal neurons (*p <* 0.0001, *KS* = 0.28; Figure 2**I**). This parameter could therefore serve as the causal mechanism behind the observed increase in ERP peak during the presence of conspecifics. Since the mean-field model used in this scale makes the inference process computationally tractable, we ran the current gold standard sampling algorithm for an asymptotically exact estimation of connectives from ERPs (Supplementary subsection 6.6). Our results indicate similar findings regarding the increased *g*_2_ during the presence of conspecifics, thus enhanced *SE_e_* (see Supplementary Figure S10). This algorithmic consistency validates our Bayesian learning using low-dimensional data features for training neural density estimators to conduct causal inference on SE. Finally, we computed the excitatory (*e*) and inhibitory (*i*) postsynaptic potentials (EPSPs/IPSPs; Figure 2**J**) by inserting the median of the pooled effective connectivity into the following equation: 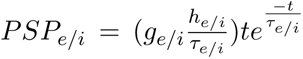. This shift in *g*_2_ increases the E/I ratio, which could, in turn, be responsible for the observed increase in peak evoked activity within the column (Figure 2**J**). While E/I ratios themselves could be non-identifiable, in our case, the similar values of IPSPs from our results in conjunction with elevated EPSPs during the presence condition alleviates this issue. Therefore, we posit that mere presence effects could be only driven by enhanced *SE_e_*. In the following, we investigate this hypothesis by performing inference from EEG recordings of a lateral-interception task at the whole-brain scale.

**Figure 2.**
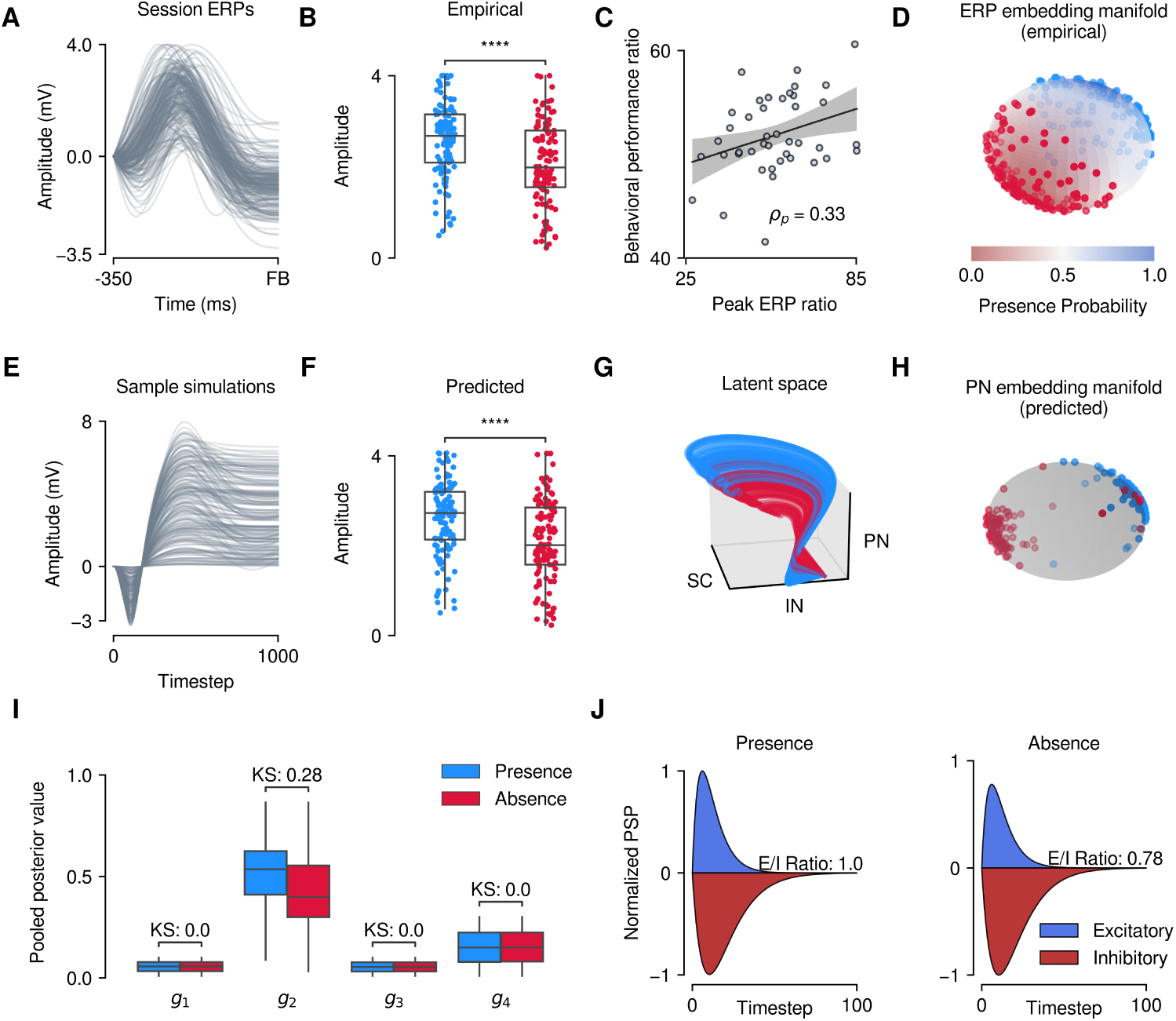
Inference of mesoscale synaptic efficacies. **A**) Feedback-locked empirical evoked potentials across all sessions. **B**) Maximum amplitude of the empirical ERPs in presence vs absence (*p <* 0.001). **C**) Correlation of between-condition ratio of ERP peak with behavioral performance (accuracy). **D**) Embedding manifold of empirical ERPs. **E**) Sample of simulations used for training the posterior density estimator. **F**) Peak amplitude of pyramidal neuron activity from posterior predictive checks across the conditions (*p <* 0.001). **G**) Nonlinear latent space of the posterior predictive check done on empirical ERPs. **H**) Embedding manifold of pyramidal neuron activity. **I**) Pooled distribution of mesoscale synaptic efficacies across sessions. **J**) Computed excitatory and inhibitory postsynaptic potentials in presence versus absence.

### 2.5. Motor responses in presence vs absence

While an increase in effective connectivity at micro/mesoscale is observed within single regions, this finding does not immediately translate to modulations of wholebrain dynamics. However, it is reasonable to hypothesize that sheer presence of conspecifics would exert disparate dynamics in functional brain networks involved in attention-modulated tasks. Not only that, the effects of social presence should theoretically manifest across a wide array of cognitive tasks [11, 39]. To this end, we designed a counter-balanced lateral interception task in which participants (*n* = 27, *n_female_* = 14, *n_male_* = 13) played a virtual game with varying ball velocities and trajectories. The task involved intercepting a downward-moving ball on a large display using a handheld slider to control a virtual paddle on-screen (Figure 3**A**). After each interception attempt, visual feedback was provided to the participant by briefly changing the paddle color to green/red for successful or failed trials, respectively. In the presence condition, the ‘observer’ sat in the participant’s peripheral vision, sometimes monitoring the hands of the ‘actor’ but not the screen, so as to minimize evaluation-induced anxiety during the presence condition [40] (Figure 3**B**). Participants were unaware of the observer’s true purpose and were informed that they were present to monitor equipment. In general, participants learned the task goal very quickly and were consistent in pursuing the ball across trials (Figure 3**C**). Kinematic and electrophysiological data (paddle movement and EEG activity) were recorded during the task (see Methods subsection 5.8). Previous studies on social facilitation have outlined the importance of using kinematic measures of performance in motor tasks, such as movement speed/duration –and features thereof– as opposed to outcome-based metrics such as accuracy or hit-rate [41]. Therefore, we analyzed average and peak paddle velocity within a 700 ms window (half-width of the longest trials) before the feedback time (Figure 3**D**). We found that average paddle speed was significantly higher in the female subject group in the presence condition compared to absence (*p <* 0.05; Figure 3**E**, left panel), whereas there was no significant difference between the experimental conditions in the male subject group (*p >* 0.05; Figure 3**E**, right panel), indicating a social facilitation effect for one subject group only.

**Figure 3.**
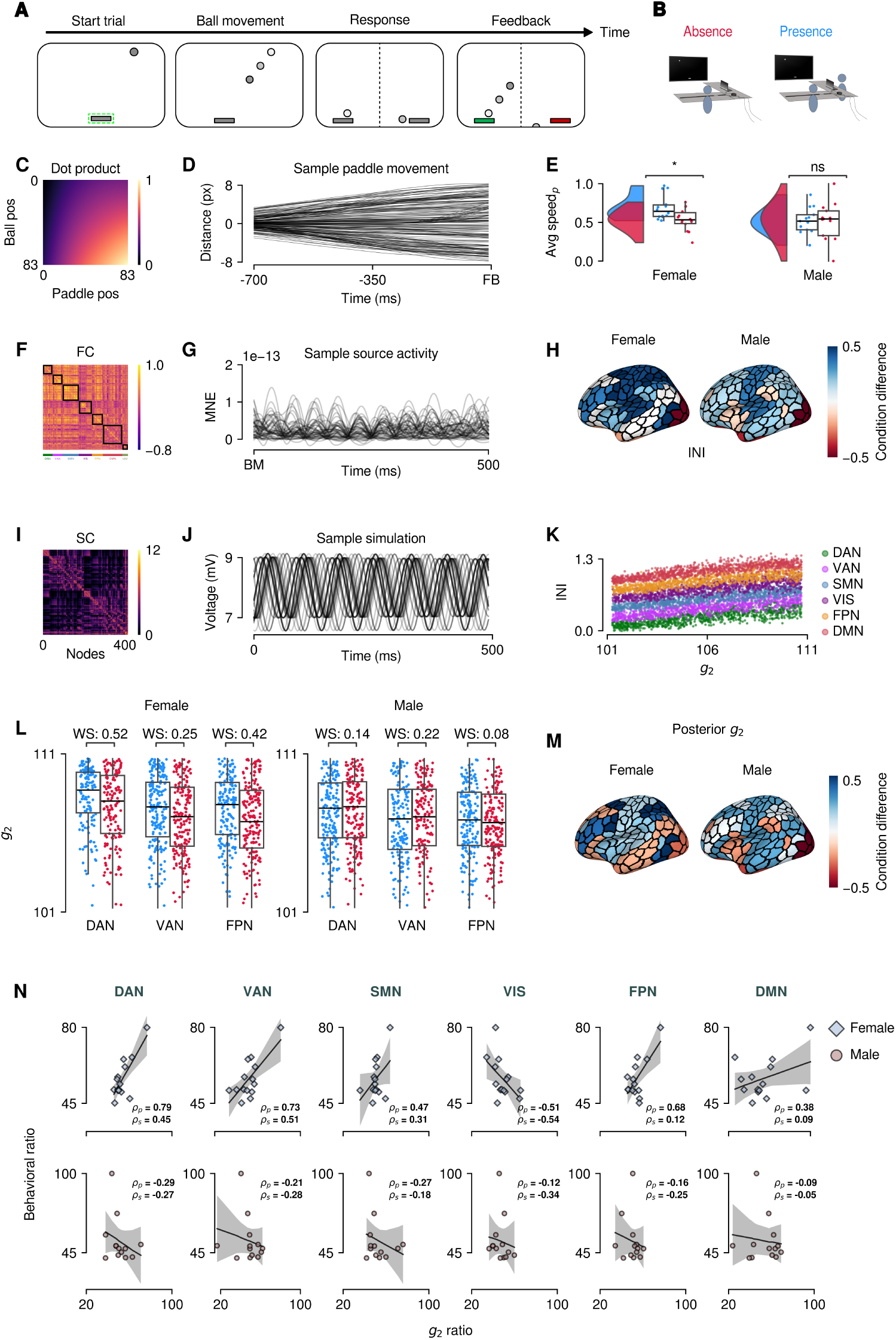
Inference of macroscale synaptic efficacy. **A**) Outline of the lateral-interception task. **B**) Schema of the lateral interception task. **C**) Covariance of ball and paddle trajectories in time, along the axis of interception. **D**) Sample paddle movement trajectories. **E**) Average paddle speed across conditions and subject groups. **F**) Sample functional connectivity from one participant, obtained by sorting according to the brain networks parcellation. **G**) Source time-courses of an example participant’s EEG. **H**) Median of the intra-network integration of the two subject groups across the seven functional networks. **I**) Average structural connectivity template used for whole-brain simulations. **J**) An example simulation from the Jansen-Rit model. **K**) Intra-network integration of random simulations used as the data feature for training the neural density estimators. **L**) Pooled estimated posterior distribution of *SEe* in dorsal and ventral attention networks across subject groups. **M**) Median of inferred *SEe* in female versus male subjects. **N**) Correlation of betweencondition ratios of inferred *SEe* versus behavioral performance (average paddle speed).

### 2.6. Increased intra-network integration during conspecific presence

Previous investigations have outlined the potential role of attention-oriented regions/networks in social facilitation [10]. If attentional modulation could indeed serve as the driving force underlying mere presence effects, one would expect to observe a higher correlation of activity in attention-oriented functional networks such as dorsal/ventral attention network, and frontoparietal network, compared to other functional networks. We thus focused solely on alpha-band activity, not only due to the prominence of alpha peak in the vast majority of the participants (see Supplementary Figure S5**C**), but also since it is prominently associated with distractor suppression, top-down attentional control, and selective attention [42, 43]. In order to investigate how attention could affect the organization of brain networks, we source-localized the participants’ EEG (Figure 3**G**) and extracted the source activity of seven functional networks [44]. These networks comprise dorsal attention network (DAN), ventral attention network (VAN), somatomotor network (SMN), visual network (VIS), frontoparietal network (FPN), default-mode network (DMN), and limbic network (LIM) (see Supplementary Figure S4). The source activity was subsequently used to compute functional connectivity (FC) for each subject (Figure 3**F**). The sum of FC within a network –henceforth referred to as intra-network integration–, could be used to quantify the overall level of integration (or interconnectedness) of brain regions within that network. A higher intra-network integration (INI) suggests a more integrated network with stronger communication across regions, and thereby could serve as a suitable ‘observable’ proxy for increased effective connectivity within functional brain networks [45]. We found that conspecific presence significantly increases intra-network integration of attentional networks such as DAN, VAN, and FPN –compared to other networks– in the female subject group (*p <* 0.001; **Figure 3H**, left). As for the male subject group, we found decreased measures of INI during conspecific presence, however, this difference was not significant (*p >* 0.05; Figure 3**H**, right).

### 2.7. Increased SE_e_ in attentional networks during conspecific presence

Intra-network integration might convey some useful information about changes to effective brain connectivity [46, 47]. However, it is not a direct measure and therefore cannot be readily relied upon for direct interpretation of changes to this connectivity. As mentioned before, we identified local recurrent excitation (the parameter *g*_2_ in the Jansen-Rit model, given by Equation 2; Figure 2) as the most significant driver between presence and absence in dlPFC and ACC. A reasonable prediction therefore entails recurrent excitation also increasing in attentional networks during conspecific presence. To test this hypothesis, we employed the framework of Bayesian learning on a whole-brain model of brain comprised of interconnected Jansen-Rit mass models (given by Equation 3, with 400 regions using Schaefer atlas; Figure 3**I**). We constrained the values of the connectivity (*g*_2_) to those corresponding to the alpha band (Figure 3**J**), and trained the neural density estimators to per-network INI values obtained from random simulations (*n* = 20000), given the linear relationship between *g*_2_ and INI values (Figure 3**K**). Empirical INIs were subsequently used to obtain joint posterior distributions of local recurrent excitation in the brain functional networks. In the female subject group, pooling the estimated posterior distributions across subjects showed a significant increase in DAN, VAN, and FPN in the presence condition compared to the absence condition (*p <* 0.001, *W S* = 0.52, *W S* = 0.25, *W S* = 0.42, respectively; Figure 3**L** left panel, Figure S6). The male subject group however, exhibited a significant decrease of SE in DAN and VAN, and a -negligible- decrease in FPN, during the presence condition (*p <* 0.001, *W S* = 0.14, *W S* = 0.22, *W S* = 0.08, respectively; Figure 3**L**, right panel). In both subject groups, the aggregate distribution of *g*_2_ across the networks showed a striking resemblance to the mapping of empirical INI values (Figure 3, panel **H** vs **M**). Since raw values of effective connectivity might not necessarily be linked to the aforementioned behavioral modulations, we calculated per-subject between-condition ratio (percent change) of average effective connectivity and that of behavioral performance (average paddle speed). The most significant finding was in the facilitated subject group (females), where we found a significantly high correlation between effective connectivity and behavior in both DAN and VAN (*ρ_P earson_* = 0.79, *ρ_Spearman_* = 0.45 for DAN, and *ρ_P earson_* = 0.73, *ρ_Spearman_* = 0.51 for VAN; Figure 3**N**). As for FPN, while we found a strong linear correlation with behavioral performance, the significance of this correlation quickly diminished after computing correlations on ranked data using spearman test (*ρ_P earson_* = 0.68, *ρ_Spearman_* = 0.12; Figure 3**N**). These results provide support for the potential role of effective connectivity in presence-induced attentional modulations. Similar to FPN, we found high linear correlations in SMN and DMN, which significantly diminished when correlations were computed on ranked datasets (e.g. *ρ_Spearman_* = (0.31, 0.09) for SMN, and DMN in the female subject group, respectively; Figure 3**N**). We found no significant correlation with behavior within any of the attentional networks in the non-facilitated group (Figure 3**N**, lower panels). Finally, we found a negative correlated relation between effective connectivity and behavioral performance in the VIS network in both subject groups. The female subject group exhibited a much stronger negative correlation of SE with behavioral performance compared to the male subject group (*ρ_Spearman_* = (−0.54, 0.34) for the female and male subject groups, respectively; Figure 3**N**). This finding further alludes to the potential contribution of attentional modulation of visual stimuli –as opposed to sheer perception of visual cues– to the observed social facilitation effects in our visuomotor task. However, we should note that due to the limited number of subjects per group, the generalizability of these findings needs to be further explored in future investigations.

## 3. Discussion

Despite the wide array of research regarding the influence of social stimuli on both brain and behavioral dynamics [48, 49, 50], the majority of this work largely revolves around some form of explicit interaction or communication between the social agents. Yet, a basic, and perhaps most fundamental component of social cognition is the awareness of other’s being present in the immediate environment, without any overt interaction. Modulations of performance during the mere presence of conspecifics have been hypothesized to largely stem from attentional processes, via the activity of context-specific neural substrates [10, 11]. However, our understanding of the neurobiological mechanisms underlying such a modulation is still severely lacking. One reason could be the difficulties in making inferences, particularly across scales. These challenges notwithstanding, recent advances in probabilistic machine learning, such as simulation-based inference (SBI; [27, 28, 51, 29, 52, 53]) have helped bridge this gap, as demonstrated in this work. We have further advanced this progress by opting for SBI that operates across brain scales. This enabled us to universally evaluate our hypothesis informed by low-dimensional features such as firing rate or level of integration. Notably, SBI bypasses the need for Monte Carlo sampling, whose gradient-based variants becomes computationally expensive when dealing with inference at the whole-brain scale, or inapplicable at the micro-scale due to the discrete nature of spike events. Moreover, this approach is amenable to an amortized strategy [28, 52], allowing the use of the same trained model, validated on synthetic data (see subsection 6.5), immediately on empirical data. From a computational perspective, this is exceedingly beneficial as it enables parallel simulations, whereas Monte Carlo is embarrassingly parallel by running different chains on different computational units [54].

In this study, we leveraged an efficient form of Bayesian learning with deep neural density estimators, to investigate the link between attentional modulation and effective connectivity –as a proxy for the activity of context-specific neural populations– during others’ presence in visual/visuomotor tasks across three spatiotemporal brain scales. In single neurons (microscale), we found that others’ presence strongly increases the overall *SE_e_* (or a decrease in the ratio of inhibitory to excitatory weights) in regions dlPFC and ACC, both of which are highly involved in attentional modulation. Our findings for the ERPs (mesoscale) further supported these results, as we observed a significant increase in local recurrent excitation in the presence condition. The observed facilitation of learning during conspecific presence supports the hypothesis of re-allocation of attentional resources due to the suppression of visual distractors. These findings are consistent with the previous research on the interplay between connectivity and attentional modulation in ACC and dlPFC. For one, stronger cooperation (i.e. the presence of significant effective connectivity) between the two regions has been reported to be critical for attention shifting [55]. In addition, alterations to the connectivity of these regions have also been linked to clinical measures of inattention and impaired cognitive control [56, 57]. Finally, the synaptic specializations in ACC suggest its potentially great impact in reducing noise in dorsolateral areas during challenging cognitive tasks, and the disruption of said mechanisms in neuropathologies such as schizophrenia and major depressive disorder (MDD) [58, 59].

Given our hypothesis, one would expect functional brain networks involved in attention to be more predictive of behavioral performance compared to others; The lateral interception task in humans complemented our findings at the preceding scales by attempting to answer this question. We found that the female subject group exhibited significantly faster average paddle movements during the presence condition compared to the absence condition, while the male subject group showed no difference in performance between the two. Similar to our findings in the micro/mesoscale, we observed an increase in effective connectivity during conspecific presence, but interestingly only in the facilitated subject group (females). Remarkably, we found that effective connectivity is strongly correlated with task performance but only in attention-oriented functional networks such as DAN and VAN. One can interpret the mutual correlation of both attentional networks with behavior as a dynamic interplay between these two networks, influenced by the task’s attentional demands [60]. This co-activation may involve the transfer of response decisions from focused targets to high-level centers [61], and is often associated with changes in synchronization and functional connectivity [62, 63]. The involvement of the DAN in attentional orienting and the modulatory influence of the VAN during reorienting [60] further support the idea of a dynamic interplay between these networks. The significant elevation of SE in the FPN during conspecific presence in the facilitated subject group, coupled with the lack of significant correlations between SE and task performance might seem initially perplexing. However, one potential explanation might stem from the lower cognitive demands of the lateral interception task compared to the association learing task. Taken together, these findings suggest that the social facilitation effect observed in females is mediated by changes in the way attentional networks coordinate and process information during presence. The observed increase in effective connectivity within these networks could reflect a more synchronized and efficient neural processing mode, ultimately leading to faster and more accurate motor performance. In sum, we suggest facilitative effects arising from others’ mere presence could largely stem from an increase in *SE_e_* (i.e., effective connectivity) in attentional brain networks, the influences of which could be observed across spatiotemporal brain scales.

### 3.1. Alpha-band coherence and attentional modulation

In our opinion alpha band FC can serve as a more reliable measure of attentional modulation than power, despite the existence of alpha peak in the majority of our subjects (see Supplementary Figure S5**C**). This approach focuses on the level coordination of activity between the nodes, which is a hallmark of focused attention [64]. Imagine an orchestral composition, the power analysis of which would provide us with a measure of ‘loudness’ of the instruments. In contrast, phase synchronization (or activity correlation) of the instruments would allow us to understand if they are playing in time, which is crucial for a cohesive melody. Indeed, alpha-band synchronization reportedly plays a crucial role in attentional modulation (particularly in the context of visuospatial attention), is reportedly modulated by the orienting of attention, and is associated with decreased reaction times to attended stimuli [42, 43]. Moreover, attention has been reported to drive the synchronization of alpha and beta rhythms between the right inferior frontal and primary sensory neocortex [65]. Finally, alpha-band phase has been shown to modulate bottom-up feature processing and is modulated by the nature of the recruited attentional mechanisms [66, 67].

### 3.2. Homeostatic framework of effective connectivity in social brain networks

The assumption of behavioral modulation observed during conspecific presence stemming from the activity of context-aware (or social) neurons, alongside our findings of altered E/I ratio during presence, leads one to expect perturbations to the functioning of said cells in social neuropathologies such as ASD and schizophrenia. Indeed, alterations in E/I balance within neural microcircuitry have been linked to these social neuropathologies [68]. Neural communication and disruptions in E/I balance in autism, stemming from atypical brain connectivity, have been linked to changes in attentional modulation and social interactions [69]. Functional connectivity between the cerebellum and cortical social brain regions is also reported to be altered in autism [70]. In schizophrenia, elevated activation of frontal control networks and association cortices has been proposed as a compensatory mechanism for impairments in connectivity within the social brain networks [71]. In addition, resting-state networks are reported to be differentially affected in schizophrenia, with reduced segregation between the default mode and executive control networks [72], as well as reduced connectivity in the dorsal attention and executive control networks [73]. Direct comparisons between the two pathologies have also found a reduction of activity in regions within the social-cognitive network in autism compared to schizophrenia [74]. It should be noted, however, that the specific nature of these changes in connectivity, the direction of said changes, and their relationship to social neuropathologies remain a topic of ongoing research [75, 76].

### 3.3. Scope and limitations

As previously mentioned, the literature surrounding the neurobiological underpinnings of the mere presence of others in higher primates is rather sparse. We attempted to address this sparsity by employing our Bayesian learning framework across three spatiotemporal brain scales– an endeavor which, to the best of our knowledge, has not been undertaken neither in the field of computational neuroscience, nor when pertaining to mere presence effects. However, there are some limitations in both of the previously outlined tasks that need to be addressed rigorously in order to obtain an accurate perspective on mere presence. The longitudinal nature of the association learning task is highly informative as it provides insights into facilitation over a much longer timescale, and partially complements our lateral interception task, but remains limited by factors such as number of subjects and sex. And considering the somewhat bold claim of ubiquity of context-aware ‘social’ neurons across the brain, future investigations in non-human primates should aim to cover a wider range of brain regions. We addressed some of these restrictions in the lateral-interception task, but there is still much to be done to refine an impeccable task outline for SFI effects. For instance, the lack of improvements of kinematic performance during conspecific presence in the male subject group might partially stem from the level of perceived rank within the immediate social environment (in this case, the task space; [77]). Thus, one should take care as to not generalize the lack of improvements during the presence condition in the male (or any) subject group as an inherent difference between the subject groups in every facilitation task. And while we set the number of participants to an acceptable threshold for between-group comparisons, another potential limitation stems from the low number of subjects in the lateral interception task. We believe that inquiries on social facilitation –by definition– require an expansive array of participants, in order to thoroughly analyze the effects of others’ presence across relevant subject groups. Finally, it is important to note that while social facilitation effects could range from extremely significant, to barely noticeable [9], such effects have been repeatedly observed in a vast array of previous investigations. Therefore, this phenomenon could currently serve as the best proxy to investigation of ‘pure’ mere presence effects. Nonetheless, the development of alternative cognitive frameworks on the underpinnings of mere presence would be invaluable for gaining a deeper understanding of the contribution of such stimuli in social cognition as a whole.

## 4. Conclusion

This study investigated the neural mechanisms underlying social facilitation, the phenomenon of improved performance in the presence of others. By conducting a multi-scale inference across monkeys and humans, we provide evidence that mere presence effects are mediated by increased effective connectivity within attentional brain networks. In monkeys performing an association learning task, the presence of an observer led to a rise in only excitatory synaptic efficacy, *SE_e_*, in the dlPFC and ACC, brain regions crucial for attention. Human participants performed a lateral interception task while being observed by a non-interacting spectator. We observed facilitated performance in one subject group, characterized by faster average paddle speeds during social presence. We were able to link this facilitation to increased *SE_e_* within the DAN and VAN, suggesting a more synchronized and efficient processing mode. These findings offer a novel perspective on social facilitation, highlighting the role of attentional networks and effective connectivity. Furthermore, the observed alterations in E/I balance suggest potential links to social neuropathologies such as autism and schizophrenia. Future research should explore the generalizability of these findings across a wider range of tasks, subject groups, and social contexts. Overall, this study sheds light on the neural underpinnings of social facilitation and paves the way for further investigation into the complex interplay between mere presence, attentional modulation, and brain network dynamics.

## 5. Materials and Methods

### 5.1. Between-Condition ratio

We defined the between-condition differences to behavioral and neural metrics *X* in all three scales, as the percent change of *X* between the presence *P* and absence *A* conditions: 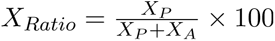.

### 5.2. Association learning: Subjects

The subjects comprised of two adult male Rhesus monkeys (Macaca mulatta). They were housed together since the age of 3 years and weighed 8-12 kg at the time of the study. They had established stable and spontaneous social interactions, with monkey *A* being the dominant as revealed by the ‘access to water and food’ test. Animal care, housing, and experimental procedures conformed to the European Directive (2010/63/UE) on the use of nonhuman primates in scientific research. The two monkeys were maintained on a dry diet, and their liquid consumption and weight were carefully monitored. While both subjects exhibited social facilitation effects, the neural recordings from Monkey *M* were highly laden with noise, and we thus opted to only use the neural/behavioral data obtained from Monkey *A*.

### 5.3. Association learning: Behavioral procedures

Monkey *A* (the actor) was trained to associate abstract images with targets on a touchscreen, either in presence of monkey *M* (the spectator) or in absence (Figure 1**A B**). Under social presence, the two monkeys were positioned in primate chairs facing each other, with their head immobilized (Supplementary subsection 6.2). Only the actor had access to the touchscreen, and thus performed the task and received rewards on correct trials. The spectator was not rewarded, had no incentive to produce any particular behavior, was never tested (as actor) during the same day, and when tested, a new set of stimuli was used (therefore preventing any observational learning from occurring). When the actor was tested in absence, the other remained in the housing room located at a distance such that the actor was truly alone in the testing box, deprived of any communicative means with the conspecific through visual, auditory, or olfactory channels. During task, the actor started trials by touching a white rectangle, which triggered the presentation of a cue at the center of the screen. The monkey was required to indicate among the four white squares (targets), the one associated with this cue. After a variable delay (500-700 ms), the cue went off (go signal) and the monkey had to move the hand and touch the chosen target. If correct, a green circle (positive feedback) informed the monkey that the choice was correct, and a reward (fruit juice) was delivered after 1 s. If the choice was incorrect, a red circle signaled the error (negative feedback), and no reward was delivered.

### 5.4. Association learning: neuronal firing rates

Social and asocial neurons were found in similar proportions in both prefrontal areas (*n* = 376 in dlPFC and *n* = 216 in ACC), and histological reconstruction revealed no spatial segregation within individual areas. Because of similar profiles and proportions of outcome-related activations in dlPFC and ACC, the data were pooled together into a single neuronal sample. The correlation of firing rates with behavior was conducted on the entire neural dataset, from which we chose a subset of neurons (*n* = 92) oriented to negative feedback –as performance in this condition correlates highly with neural firing rate– which belonged to Monkey A and were strictly categorized as social or asocial. A time window of 500 ms before and after the feedback time was used. Since we were interested in the difference of firing rate between the conditions per neuron, the neural firing rates were normalized by dividing by the maximum firing rate of each neuron.

### 5.5. Association learning: event-related potentials

Local-field potentials (LFPs) across four channels were epoched by time-locking the signals to a 350 ms time window before the feedback timestamps. A high-pass filter of 2 Hz was applied to the LFPs before epoching, and electrodes with flat signals were removed from the analysis. The electrodes were further filtered by including only those with excellent and good signal quality in the calculation of ERPs. Trials with negative feedback were averaged during the presence or absence to obtain the raw ERPs (*n* = 117) in each task condition. A low-pass (butterworth) filter of 20 Hz was then applied to the evoked potentials, which were then up-sampled to 1000 time-steps. Subsequently, evoked potentials in each conditions were normalized with respect to the maximum amplitude of their session in order to retain across-session comparability. These normalized evoked potentials were then used as input to the machine learning method CEBRA [78] to obtain low-dimensional embeddings of the data. Using the PyMC [79, 80] package, the embeddings themselves were fed to a Bayesian neural network using variational inference with a perceptron network with two hidden layers and 5 nodes per layer. This process aimed to obtain a smooth probability grid of the likelihood of an ERP embedding belonging to one of the experimental conditions.

### 5.6. Lateral interception: participants

A total of 28 healthy right-handed participants (14 women and 14 men, with an average age of 23.6 ± 2.1 years) were recruited for the study. All participants provided written consent before participating in the study. One male participant was removed from the study due to a recording error (*n_male_* = 13). The study procedures were approved by the Central Ethics Board of the University Medical Centre of Groningen (UMCG) and according to the Declaration of Helsinki (World Medical Association).

### 5.7. Lateral interception: experimental setup

In the presence condition, a ‘spectator’ sat across from the participant in their peripheral vision at a distance of 1.5 meters, meaning that the spectator sat in the participants’ social space, as defined by Hall [81]. The spectator’s sex and age were controlled to be close to those of the participants, and participants were unaware of the true purpose of the spectator’s presence, as they were informed through a cover story that the spectator was present to monitor the equipment, with the spectator appearing to focus on a laptop. To prevent the spectator from inducing a sense of evaluation, they avoided looking at the screen and focused approximately 70% of the time on observing the participants’ hands, thereby creating a sense of being observed.

### 5.8. Lateral interception: data preprocessing

Behavioral data were epoched to a 700 ms time window preceding the feedback events. This duration was long enough to encompass the majority of paddle movements, but also allowed for the preclusion of the starting flat tails of paddle movements. A low-pass 5 Hz filter (*order* = 2) was subsequently applied to the movement data. Paddle data for each subject were then normalized with respect to ball arrival positions and subsequently converted to absolute values to ensure uniform computation of paddle speeds. In addition, artefactual trials (with misaligned starting position) were excluded from the analysis.

Preprocessing of whole-brain recordings was conducted automatically and agnostically, minimizing biases in preprocessing pipeline due to experimenter’s subjective choices–, and was entirely performed using the MNE-python package [82, 83, 84]. First, we identified and interpolated bad EEG channels. This was done (per-subject) by creating trials from -500 ms to 1000 ms after ball movement (with a baseline of -500 ms to -200 ms), which were given to the Ransac module of the AutoReject package [85]. EEG channels were then re-referenced using average referencing. Second, we removed blinks and saccades components from EEG via independent component analysis (ICA). EEG data were first filtered using an FIR band-pass filter from 1 to 45 Hz (using the ‘fastica’ method), of which, epochs of equal lengths alongside a rejection threshold were computed, which were then given to the ICA algorithm for computation of the independent components. General templates for blink and saccade components were initially computed and saved from a single subject (*p*01). Subsequently, these components were template-matched and projected out of the data (*threshold* = 0.9) for all participants. Due to the portable nature of the EEG setup, which was specifically designed for movement in the natural environment, we did not have access to subject-specific MRI scans. Therefore, source-localiztion was performed via the freesurfer [86] template ‘fsaverage’ (for comparison between experimental conditions, the data need to be morphed to a representative ‘template subject’ regardless, even in the presence of per-subject structural imaging). The source space model ‘fsaverage-ico-5-src’) and boundry-element model (‘fsaverage-5120-5120-5120-bem-sol’) were used to compute the general forward solution (*mindist* = 5*mm*). Noise covariance was computed from broadband epochs spanning from -500 ms before to 1000 ms after ball movement, and a baseline of -500 ms to -200 ms (with a *tmax* = −0.2 ms). The forward solution and the noise covariance were then used to create the inverse operator (with regularization parameter *loose* = 0.2).

Extraction of source time-courses is as follows. The data were first band-passed (multi-taper method) to the alpha band (8-12 Hz), and subsequently epoched for a time window of 500 ms starting from ball movement (with a baseline from -200 to 0, cropped before saving the epochs). We chose this window as attentional modulation should theoretically occur before modulations to behavior, and closer to the visual cue. Subsequently, the precomputed inverse operator was used with the MNE algorithm to obtain source activity in vertex format, which was in turn used alongside the Schaefer 2018 atlas (400 parcels) to extract the source time-courses for the activity of the functional brain networks [44, 87].

### 5.9. Lateral interception: functional connectivity

Functional connectivity (FC) was calculated as the Pearson correlation between the sources/labels (or nodes in case of simulations). For the correlation matrix FC, the element *FC_ij_* representing the statistical dependency between two labels *i* and *j*, was calculated as 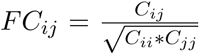, where *C_ij_* is defined as the covariance between said labels, and *C_ii_* as the variance of *i* [18]. Elements of the FC matrix were subsequently sorted and isolated into separate masks according to the aforementioned functional networks (with left and right hemispheres concatenated together, as we are interested in the activity of the entire network). The sum of their upper triangular was used to obtain intra-network integration: 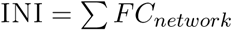.

Our choice for this coherence measure is supported by previous research, as not only the strength of distractor suppression is reportedly influenced by the intrinsic connectivity within and between attentional networks, but within-network connectivity was found to be a better predictor of this suppression [88].

### 5.10. Generative Model of Spiking Neurons

The model employed at microscale is a network of spiking neurons with alpha synapses, comprising two subpopulations: excitatory (*n* = 10000) and inhibitory (*n* = 2500) leaky integrate-and-fire (LIF) neurons [36]:

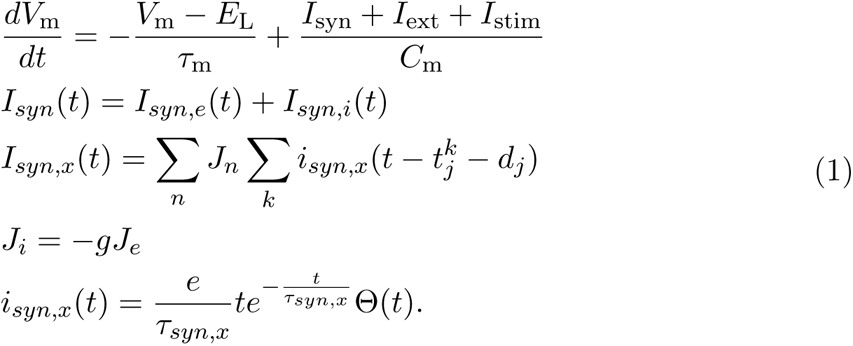

If *V*_m_(*t_k_*) *< V_th_* and *V*_m_(*t_k_*_+1_) ≥ *V*_th_, then a spike is emitted at timestep *t*^∗^ = *t_k_*_+1_ and the membrane potential is reset to *V*_m_(*t*) = *V*_reset_ for *t*^∗^ ≤ *t < t*^∗^ + *t*_ref_.

With regards to the synaptic currents *I_syn,x_*, subscript *x* represents either excitatory (*e*) or inhibitory (*i*) synapses, and both synaptic subtypes share an identical time constant *τ_syn_*. Here, *n* serves as the index for neurons within either the inhibitory or excitatory subpopulation. For a neuron with index *n*, then *k* and *d_j_* represent the index for spike times of, and the delay from said neuron, respectively. The remaining parameters and their representations are as follows:

Moreover, the external current *I_ext_* is modeled as a Poisson generator with a rate of *p_rate_* = 13341.8 *Hz*, which is calculated based on the in-degree of the excitatory synapses, the external rate relative to threshold rate, and a number of other parameters (see [89] and [36] for in-depth discussion of the external current). In order to simulate the effects of feedback anticipation in the population, a step current *I_stim_* from 350 *ms* to 900 *ms* was applied to the population of neurons in each simulation, with mean of 150 *pA* and standard deviation of 1 *pA*. During each simulation neurons received a different realization of the current based on the standard deviation of the current: *I_stim_*(*t*) = *µ* + *σN_w_*, where *N_w_* is a sample drawn from the zero-mean unit-variance normal distribution for each time-step during the current’s activation interval *w*.

We employed the NEST simulator [89] and simulated the model (*n* = 10000) for 1000 ms (*dt* = 0.1 *ms*) with varying values of the parameter *g*, which represents inhibitory/excitatory weight ratio. For each simulation, the value of *g* was randomly sampled from a truncated prior uniform distribution *g* ∈ [5, 8]. All other parameters of including external input, leak time constant, and synaptic current time constants were fixed (Table 1). Consequently, simulation firing rates were computed by first calculating the histogram of the spike times (*nbins* = 100), which were then smoothed with a Butterworth low-pass filter of *order* = 5 and critical frequency of 2 Hz (20 Hz in the original sampling rate). Firing rate time series were then normalized between 0 to 1, and the maximum value of each time series was computed. After training the deep neural density estimators (see Methods subsection 5.14), empirical firing rates were used as low-dimensional data features to efficiently obtain the posterior distributions of effective connectivity given each condition and neuron. The training took approximately two minutes, while the sampling took less than a minute.

**Table 1.** Parameters for LIF neurons.

| Parameter | Representation | Value |
| --- | --- | --- |
| $\tau_{syn}$ | Synaptic time constant | 0.5 *ms* |
| $\tau_m$ | Membrane time constant | 20.0 *ms* |
| $\tau_{ref}$ | Duration of refractory period | 2.0 *ms* |
| $C_m$ | Membrane capacitance | 250.0 *pF* |
| $V_{th}$ | Spike threshold | 20.0 *mV* |
| $E_L$ | Resting membrane potential | 0.0 *mV* |
| $V_r$ | Reset potential of the membrane | 10.0 *mV* |
| $V_m$ | Membrane potential | 0.0 *mV* |

### 5.11. Generative Model of ERPs

The generative model of ERPs used for the inference at the single column (mesocspoic level) is based on a modified iteration of the Jansen-Rit neural mass model (NMM; [90]) developed for dynamical causal modeling (DCM; [17]). The model comprises ten parameters as *g*_1,2,3,4_ (connection strengths), *τ_e/i_* (membrane rateconstants), *h_e/i_* (post-synaptic potential maximum amplitude), *δ* (intrinsic delay), and *u* (external input). The NMM model comprises nine ordinary differential equations of hidden neuronal states *x*(*t*) that are a first-order approximation to delay-differential equations, i.e., using *x*(*t − δ*) = *x*(*t*) *δẋ*(*t*). The dynamics of three neural populations: pyramidal neurons *x*_9_, spiny-stellate cells *x*_1_, and inhibitory interneurons *x*_7_ are given by:

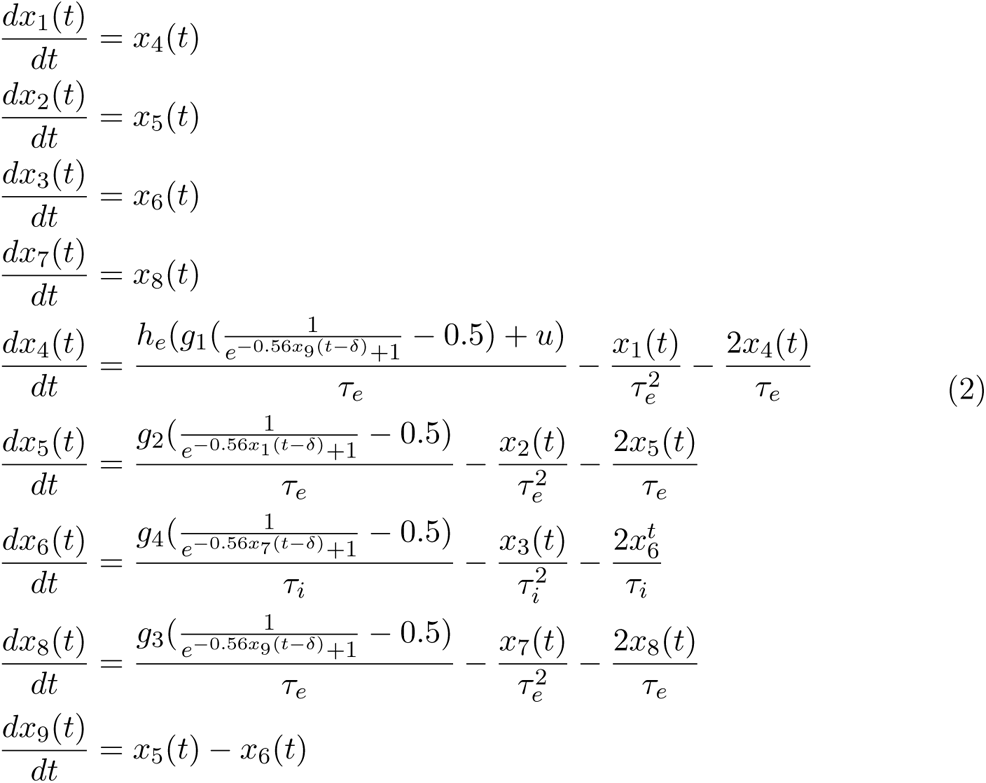

The fixed parameters for the simulations are given in Table 2. For the inference at mesocspoic scale, the samples were drawn from a uniform random distributions with minima and maxima set given in Table 3.

**Table 2.** Parameters for ERP model.

| Parameter | Representation | Value |
| --- | --- | --- |
| $\delta$ | Intrinsic delay | 8.41 |
| $\tau_i$ | Inhibitory membrane rate constant | 7.77 |
| $h_i$ | Inhibitory maximum PSP amplitude | 27.87 |
| $\tau_e$ | Excitatory membrane rate constant | 5.77 |
| $h_e$ | Excitatory maximum PSP amplitude | 1.63 |
| $u$ | External input strength | 3.94 |

**Table 3.** Prior range for ERP model parameters.

| Parameter | Representation | Min | Max |
| --- | --- | --- | --- |
| $g_1$ | Average number of synapses from PNs to SCs | 0.01 | 0.1 |
| $g_2$ | Average number of synapses from SCs to PNs | 0.02 | 1.5 |
| $g_3$ | Average number of synapses from PNs to INs | 0.01 | 0.1 |
| $g_4$ | Average number of synapses from INs to PNs | 0.01 | 0.3 |

### 5.12. Generative Model of Whole-brain EEG

Taking a network-based approach and connectome-based modeling [91, 92, 93], we generated whole-brain EEG simulations by placing the Jansen-Rit neural mass models at each parcelled brain region, connected through a structural connectivity (SC) matrix:

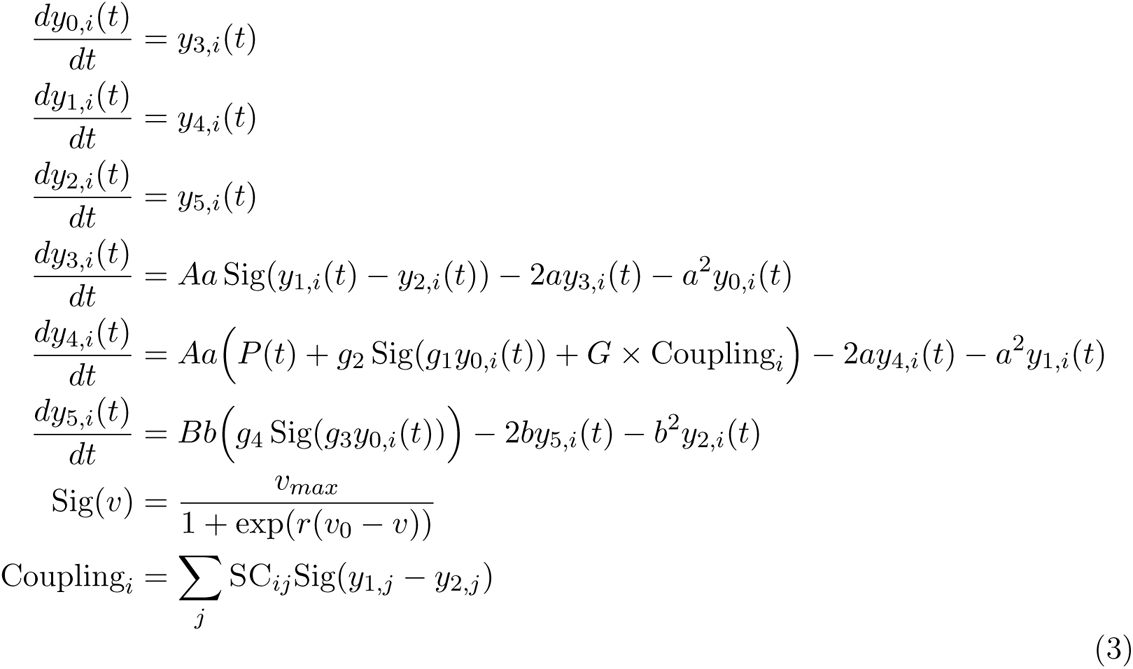

The SC matrix was obtained using tractography techniques [94], with Schaefer Atlas (400 regions), averaged across subjects, and subsequently normalizing by *SC_norm_* = log (*SC* + 1). The external current *P*(*t*) was modeled as a Gaussian random noise with mean 0.295 and zero standard deviation. All simulations were done using an Euler integration method with *dt* = 0.05 *sec*. The fixed model parameters are given in Table 4, whereas we inferred *g*_2_ (i.e., average number of synapses from SCs to PNs). The prior distribution for this parameter was defined as a truncated uniform distribution *g*_2_ ∈ [101.25, 110.7]. This range was selected to induce alpha-band oscillations in all simulations.

**Table 4.**
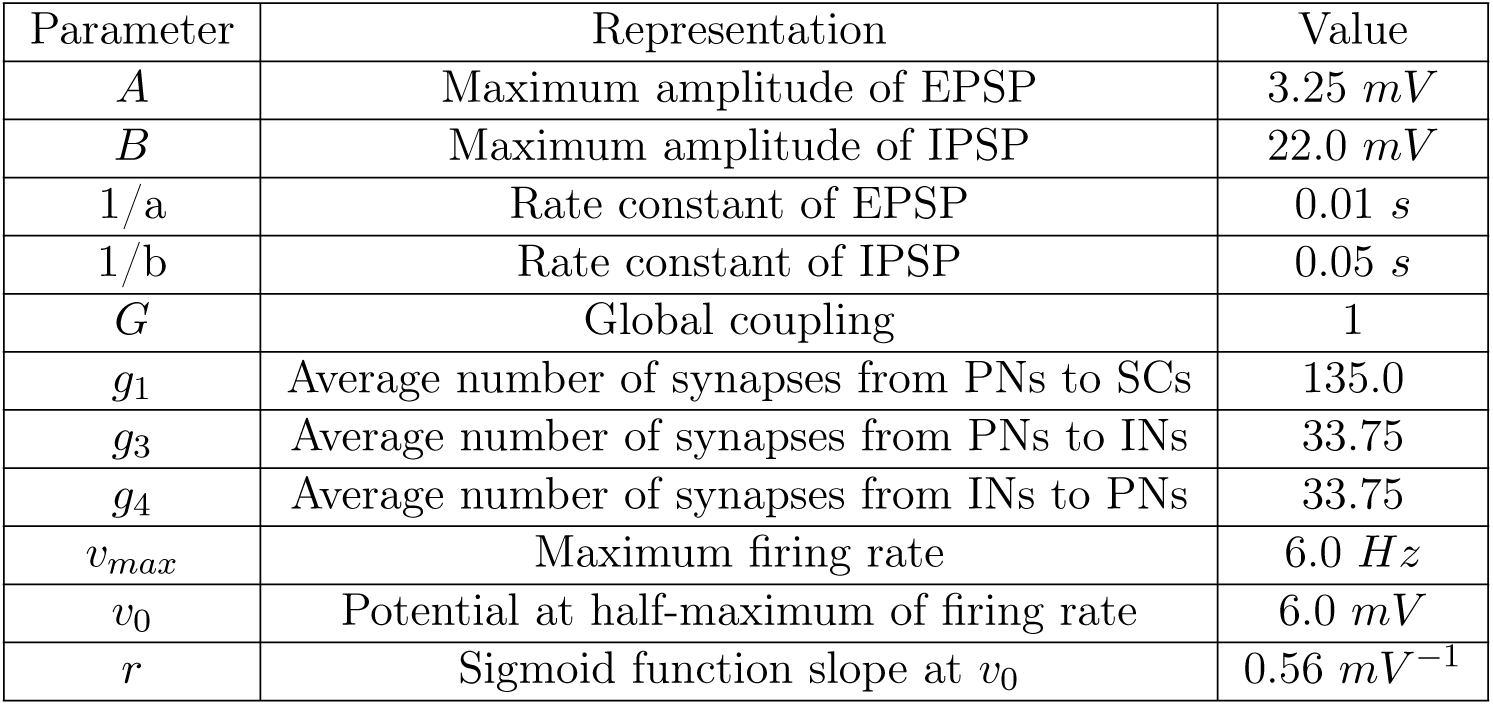
Parameters for whole-brain EEG model.

**Table 5.**
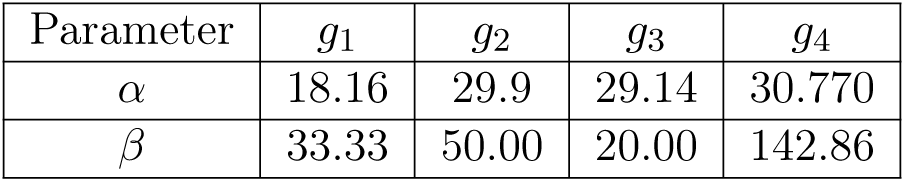
Prior distribution of synaptic efficacies in HMC.

| Parameter | $g_1$ | $g_2$ | $g_3$ | $g_4$ |
| --- | --- | --- | --- | --- |
| $\alpha$ | 18.16 | 29.9 | 29.14 | 30.770 |
| $\beta$ | 33.33 | 50.00 | 20.00 | 142.86 |

Specifically, we simulated the model (*n* = 20000 simulations) for 3000 ms (the first 2000 ms of which were cropped as transients). Simulations were then used to compute functional connectivity and intra-network integration (Methods subsection 5.9). Sampled parameters from the prior alongside INIs –as data features– were then used to train the posterior density estimators (Methods subsection 5.14). Empirical INI values were then used to sample from the learned posterior distributions of effective connectivity for each brain subnetwork.

### 5.13. Linear Dynamical Systems (LDS)

Linear dynamical systems (LDS; [95]) are widely used to model data whose evolution functions over time are linear, making them mathematically tractable for learning and inference. At time-step *t*, we observe a high-dimensional emission *y_t_ ∈* ℝ*^N^*, which is driven by low-dimensional hidden states *x_t_* ∈ **ℝ*^D^***. The state dynamics obey the equation *x_t_* = *Ax_t−_*_1_ + *V u_t_* + *b* + *w_t_*, where *A* is the dynamics matrix, *V* is the (the input-to-state) control matrix, *u_t_* is a control input, *b* is an offset vector, and *w ∼* (0, Σ*_w_*) denotes the process noise. To fully specify an LDS, we also need to describe how the emissions *y_t_* are generated from the hidden states *x_t_*. A simple linear-Gaussian emission model is given by *y_t_* = *Cx_t_* + *Fu_t_* + *d* + *v_t_*, where *C* is the measurement matrix, *F* is the feed-through (input-to-emission) matrix, *d* is an offset or bias term, and *v ∼ N*(0, Σ*_v_*) denotes the observation noise.

We used the open-source state space modeling package (*ssm*; [96]) for parameter estimation in LDS, streamlining the estimation of hidden states and model parameters from observed time series data (using Expectation-Maximisation algorithm). An LDS was constructed and fitted to the firing rate time series in each condition (*n* = 92), assuming Gaussian dynamics and emission, to learn the vector fields driven by the measurement matrix *A*, for each condition. For stability, all eigenvalues of *A* must lie strictly within the unit circle in the complex plane, which means their magnitudes must be less than 1. The fitted systems were subsequently sampled (*n* = 5000) to forecast the system dynamics for each condition.

### 5.14. Simulation-based Inference (SBI)

We adopted a Bayesian framework for likelihood-free inference and uncertainty quantification of effective connectivity at micro, meso, and macro scales. Across different brain scales, the likelihood function (i.e., the conditional probability of obtaining data given parameters) can become computationally prohibitive, rendering Monte Carlo estimation of posterior (i.e., the conditional probability of obtaining parameters given data) inapplicable. This is due to either the high-dimensional nature of the data (e.g., the large number of neurons or brain regions), or the complex relationships between variables and parameters (e.g. nonlinearity, and numerous measures of effective connectivity), leading us to employ simulation-based inference (SBI; [27]) leveraged by deep neural density estimators [28, 29]. Taking prior distribution 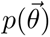 over the parameters of interest 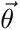, a limited number of *N* simulations are generated for training step as {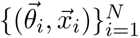, where 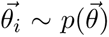 and 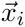 is the simulated data features given model parameters 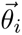*_i_*. After the training the generative models of probability distributions, so-called normalizing flows [31], we are able to efficiently estimate the approximated posterior 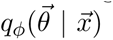 with learnable parameters *ϕ*, so that for the observed data features 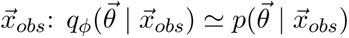.

Through a series of invertible transformations, implemented by deep neural networks, normalizing flows convert a simple initial distribution (uniform prior) into any complex target distribution (multimodal posterior). The state-of-the art neural spline flows (NSFs; [97]) offer efficient and exact density evaluation and sampling from the joint distribution of high-dimensional random variables with low-cost computation. NSFs leverage splines as a coupling function, enhancing the flexibility of the transformations while retaining exact invertibility. To conduct SBI, we used an NSF model consists of 5 flow transforms, two residual blocks of 50 hidden units each, ReLU nonlinearity, and 10 spline bins, all as implemented in the public *sbi* toolbox [98]. By training NSFs on the spiking network, neural mass model of ERPs, and whole-brain network model of EEG, we were able to readily estimate the approximate posterior of effective connectivity from low-dimensional data features, such as maximum firing rates, peak ERP amplitude, and intra-network integration. For validation of SBI on synthetic data, across brain scales see Supplementary subsection 6.5.

## Data and code availability

All computations were run on a linux machine with an AMD Ryzen 9 5900HX CPU, RTX 3070Ti GPU, and 32 GBs of RAM. All code will be available on a private github repository during the review process and all data will be available on zenodo after publication.

## Acknowledgements

This project has received funding from the European Union’s Horizon 2020 research and innovation programme under the Marie Skłodowska-Curie grant agreement No 956003. The non-human primate study was supported most notably by a National Research Agency (ANR) grant ANR-01–‘NEURO-Oox’ to D.B.; by a doctoral fellowship from Aix-Marseille University (‘bourse pre’sidentielle’) to M.D., under the supervision of D.B. and P.H., by the ‘Fondation pour la Recherche Medicale’ (FDT20130928424), and by the ‘Institut des Sciences Biologiques’ (INSB) of the CNRS to P.H. and M.D. F.I. was supported by a BDI-PED fellowship from the CNRS. V.J., and M.H. have received funding from EU’s Horizon 2020 Framework Programme for Research and Innovation under the Specific Grant Agreements No. 101147319 (EBRAINS 2.0 Project), No. 101137289 (Virtual Brain Twin Project), and government grant managed by the Agence Nationale de la Recherche reference ANR-22-PESN-0012 (France 2030 program). The funders had no role in study design, data collection and analysis, decision to publish, or preparation of the manuscript.

## Author contributions

Conceptualization: A.E., M.D., P.H., F.Z., V.J., D.B., M.H. Methodology: A.E., M.D., M.V., F.I., P.H., F.Z., A.Z., V.J., D.B., M.H. Software: A.E., A.Z., M.H. Investigation: A.E., M.D., F.I., M.H. Visualization: A.E. Supervision: P.H., F.Z., V.J., D.B., M.H. Funding acquisition: P.H., F.Z., V.J., D.B., M.H. Writing – original draft: A.E., M.H. Writing - review & editing: A.E., M.D., M.V., A.Z., F.I., P.H., F.Z., V.J., D.B., M.H.

## 6. Supplementary

### 6.1. Association learning: Behavioral data

During each testing sessions, which occurred once per day, monkey *A* learned a set of two associations under the absence condition, and an equivalent set under in the presence of monkey *M*. Each condition block consisted of 80 trials. We employed this format first and foremost due to the subject’s reluctance to performing the task in the absence block if it preceded the presence block, potentially due to anxiolytic effects of familiar conspecific’s presence with regards to learning [99, 7, 100]. In addition, previous research has alluded to lingering effects of others’ presence whilst in the absence condition, thus complicating interpretation of results in the former condition [101]. We thus chose a test-retest framework as our control, where monkeys perform the task twice in the absence condition.

The behavioral data consisted of a total of 181 sessions for monkey *A*. In order to assess the influence of mere presence on learning speed –i.e., trials-to-criterion (TTC)–, we set the sessions as the unit of analysis. TTC was defined as the third of five consecutive correct trials. We observed that mere presence reduced the number of TTC compared to the absence condition (from 11.02 to 9.26 across sessions in monkey *A*, *F*(1, 180) = 10.84; *p <* 0.001, 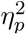 = 0.06) but a null effect of task repetition for test-retest control sessions. This indicates a social facilitation effect, which we further validated using the non-parametric Mann-Whitney U-test (*ps <* 0.05).

### 6.2. Association learning: Surgery and recording

Monkeys underwent surgery for implantation of a recording chamber over the hemisphere contralateral to the hand used for the task, along with a bolt for head immobilization. Pre-operative magnetic resonance imaging (MRI) scans of each monkey’s brain were used for chamber placement, which chamber provided access to the dorsolateral prefrontal cortex (dlPFC) and anterior cingulate cortex (ACC). A focal craniotomy was performed in the area covered by the chamber, which was sealed with a removable plastic cap during recording sessions.

We utilized a multi-channel recording system (Alpha Omega Alphalab) to acquire extracellular neuronal signals from the brain. Single tungsten microelectrodes (impedance 0.8 − 1.2 *M*Ω, FHC Instrument) were advanced into the targeted cortical regions, the dlPFC and ACC, guided by pre-operative MRI data. During each recording session, up to four electrodes were employed simultaneously, with two in the dlPFC and two in the ACC. The precise locations of the electrodes within each area were varied across sessions to sample extensively from the neuronal populations (See Supplementary Figure S1**B**). The raw signals captured by the electrodes underwent analog processing, including high-pass filtering at 6 kHz, low-pass filtering at 250 Hz, and amplification via the Alphalab software suite. These neuronal signals were temporally aligned and synchronized with key task events, such as the onset and offset of visual stimuli, behavioral responses executed by the animal, and the delivery of fluid rewards. Timing information for these events was acquired from the Cortex software [102]. The processed analog signals were stored for subsequent offline analysis. For spike sorting and absence of individual neuronal waveforms, we utilized a custom MATLAB toolbox in conjunction with the MClust Spike Sorting Toolbox [103]. This procedure aimed to disambiguate the action potentials emanating from distinct neurons, separating their activity patterns from background noise and the spiking output of neighboring cells. The isolated spike clusters, now representing the firing of putative single units, were further processed within the Neuroexplorer software to categorize the neurons based on their response profiles to various task events and stimuli.

**Figure S1.**
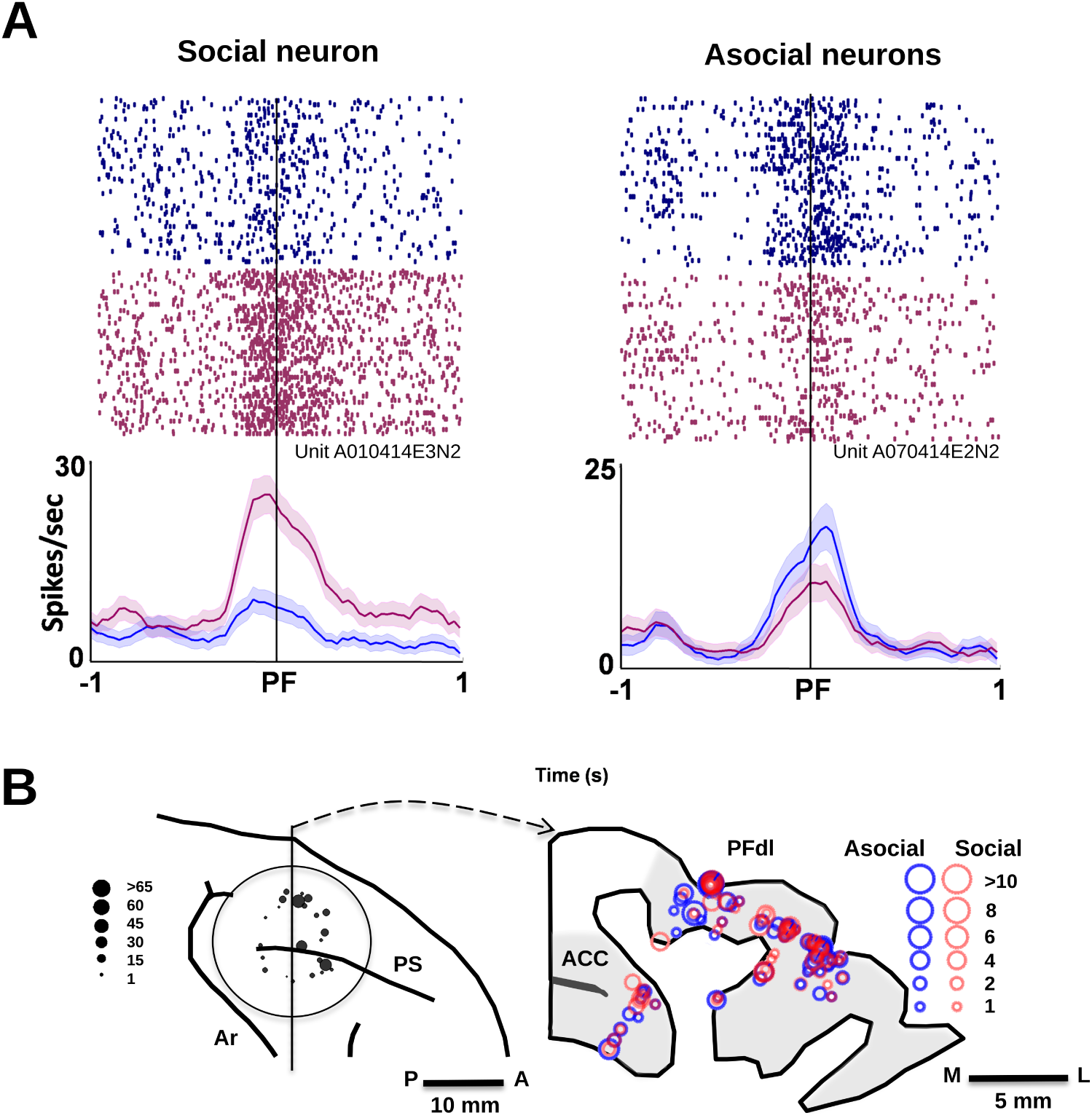
Activity and distribution of context-oriented neurons in the association learning task, during presence (in purple), and absence (in blue) conditions. **A**) Rasterplot of activity of feedback oriented (social/asocial) neurons (top), and their respective firing rates (bottom) in both experimental conditions (Presence in purple, absence in blue). **B**) Sub-areas sampled in dlPFC and ACC across the experimental sessions. Figure was adapted from our previous study [10].

For the primary analysis, we focused on the neuronal firing patterns time-locked to the onset of feedback delivery (Supplementary S1**A**). To enable statistical comparisons across conditions, we normalized the spike data by computing z-scores in 10 millisecond time bins. The z-score transformation was calculated using the following formula:

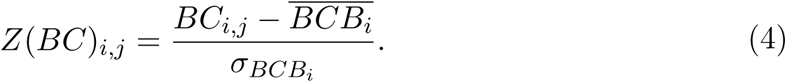

Here, *BC* and *BCB_i_* represent the bin-counts and their baselines, *i* the trial number, and *j* the bin number. T<u>herefor</u>e, *BC_i,j_* is the number of spikes in bin *j* of trial *i* during the epoch of interest, 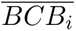 is the average baseline firing rate during the fixation period for that trial, and *σ_BCBi_* is the standard deviation of the baseline firing rates. This normalization procedure expressed the spike counts in each bin relative to the baseline activity on a trial-by-trial basis, yielding z-scored spike rasters. We then constructed peri-stimulus time histograms (PSTHs) from these normalized rasters to visualize the neuronal response profiles. The PSTHs were analyzed to categorize the neurons based on three key parameters of their discharge patterns in relation to the feedback signal: response latency, amplitude, and duration. A neuron was classified as feedback-related if its PSTH exceeded a z-score of 1.96 (95% confidence interval) for at least three consecutive bins. The latency was defined as the first bin where this threshold was crossed, while the response termination was marked by the first of three consecutive bins with *Z <* 1.96. Across the two task conditions (absence and presence), we compared the neuronal activity within each analysis epoch (See Supplementary Figure S2**A**, **B**) using Mann-Whitney U tests (*p <* 0.05). This allowed us to determine if individual neurons exhibited preferential firing in one condition over the other during the same temporal epoch.

**Figure S2.**
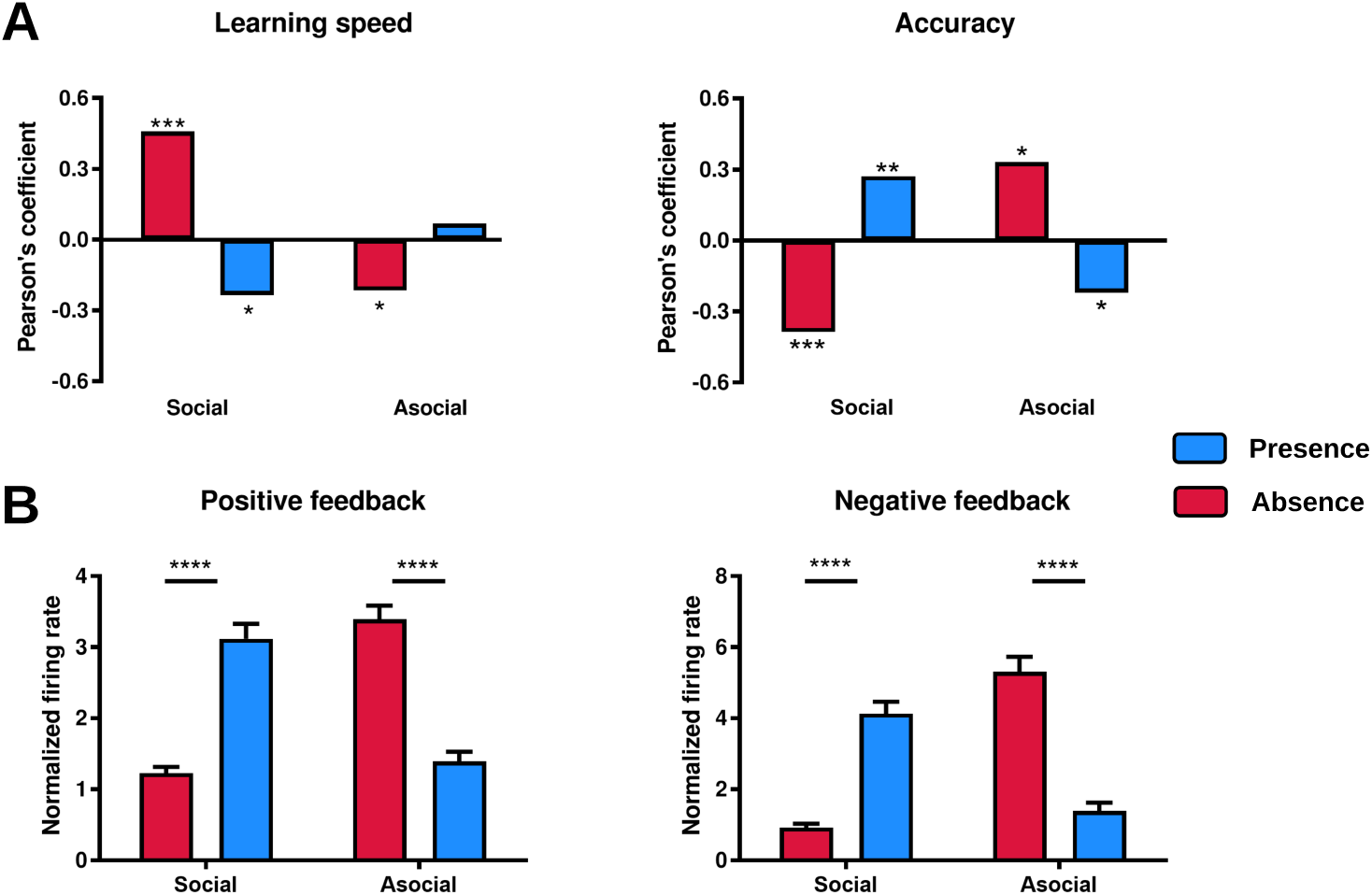
Differences of behavioral and neural patterns in presence versus absence. **A**) Correlation between the firing rate of context-specific (social/asocial) neurons and task performance. **B**) Difference of firing rate of the same neurons between the presence and absence conditions. Figure was generated using methods and data from our previous study [10].

### 6.3. Association learning: context-oriented neurons

The presence of a spectator modulated the outcome-related activity of a substantial majority of prefrontal neurons, whether they encoded errors (85%) or rewards (82%). Through our analysis, we identified two principal categories of neurons based on how their firing rate amplitudes were influenced by social presence. This categorization relied on Mann-Whitney U tests comparing the PSTHs across conditions. The first category, termed ‘social neurons’, exhibited increased firing rates when the social partner was present compared to when the animal was alone. These neurons constituted 41% and 38% of the populations responding to negative and positive feedback, respectively. The second category, labeled ‘asocial neurons’, displayed the opposite pattern, with higher firing rates during the absence condition relative to social presence. Asocial neurons represented 41% and 47% of the negative and positive feedback-related activations, respectively. To validate our categorization of neurons into social and asocial groups, we performed an unsupervised hierarchical clustering analysis on the data. This analysis corroborated the existence of two primary neuronal categories, corresponding closely (with less than 3% overlap) to the social and asocial groups identified through the Mann-Whitney U tests. Some finer subclustering was also observed within each main category. Importantly, we found that social and asocial neurons were distributed in similar proportions across the two prefrontal regions under investigation, the dlPFC, and ACC. Furthermore, histological reconstructions revealed no apparent spatial segregation of these neuronal categories within either cortical area.

### 6.4. Lateral interception: EEG experimental setup

The participants were stationed at a table positioned 2 meters away from a large 55-inch Samsung LED television screen with a resolution of 1920 × 1080 pixels and a refresh rate of 120 Hz. They were provided with a handheld knob connected to an in-house developed slider, which allowed them to laterally control a virtual paddle displayed on the screen. The objective was to intercept a virtual ball that descended vertically across the screen. Supplementary Figure S3 presents a schematic overview of the experimental setup. The positions of the slider were continuously recorded using a National Instruments data acquisition card. Kinematic data pertaining to the movements of the virtual paddle and ball were collected, along with information regarding whether the ball was successfully intercepted or not. The experiment was built and controlled using the PsychoPy software platform [104]. Concurrent with the behavioral task, electroencephalographic (EEG) signals were recorded using a TMSi SAGA data recorder and a 64-electrode TMSi infinity gel headcap (TMSi, Enschede, The Netherlands). The ground electrode was placed on the left wrist of the participant, and an online average reference was employed. To ensure consistent cortical recordings across participants, skull measurements were taken, and the Cz electrode was positioned at the intersection of the midline between the inion and nasion, and the midline between the tragus of each ear. Electrode impedances were maintained below 10 *k*Ω, and shielded cables (TMSi) were utilized to minimize artifacts arising from cable movements. The EEG signals were sampled at a frequency of 2048 Hz. Furthermore, an in-house developed software system was integrated to send triggers to the EEG docking station, marking the onset of stimuli and the precise moment when the ball crossed the invisible interception axis or made contact with the paddle. Participants were randomly assigned to one of two counterbalanced groups and engaged in a game where they had to intercept a virtual ball (ball radius = 16 *px*) descending on the screen using a virtual paddle (width = 48 *px*, height = 13 *px*). The mapping was such that 16 *px* corresponded to 1 cm on the screen. Supplementary Figure 1B illustrates the experimental design employed in the present study. To initiate a trial, participants had to position their virtual paddle within a red box located at the center of the screen. After maintaining their paddle inside the box for 100 frames (less than 1 second), the box turned green, signaling the start of the trial and the appearance of the ball. The ball velocities and trajectories were manipulated to ensure that participants achieved an intended success rate ranging between 80% and 85% successful interceptions. The ball velocities were configured such that the balls could descend the screen in either 1.4 seconds, 1.2 seconds, or 0.6 seconds. The ball trajectories were constructed in a way that allowed the balls to depart from one of five ball departure positions (BDP) along the X-axis: -672, -336, 0, 336, 672, with a fixed Y-coordinate of 501. Similarly, the balls could arrive at one of five ball arrival positions (BAP) along the X-axis: -672, -336, 0, 336, 672, with a fixed Y-coordinate of -508. To prevent any effects of familiarization, a random offset was added to the BDP and BAP on each trial. Participants first completed a 10-trial practice block to familiarize themselves with the task, followed by two 80-trial experimental blocks. During the practice block, participants encountered only trajectories of 1.4 seconds and 1.2 seconds. In the experimental blocks, participants faced 36 trajectories of 1.4 seconds and 1.2 seconds, as well as 8 fast balls (0.6 seconds) that were often un-interceptable, to maintain interest in the task and their attention levels high (Supplementary Figure S3).

**Figure S3.**
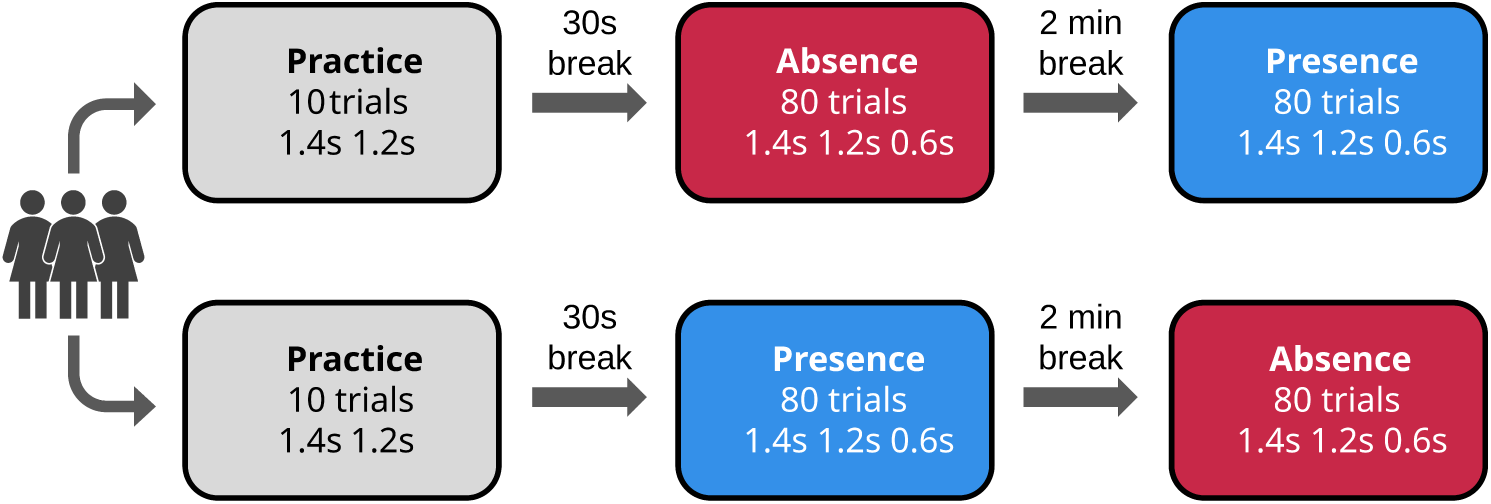
Block design of the lateral-interception task.

**Figure S4.**
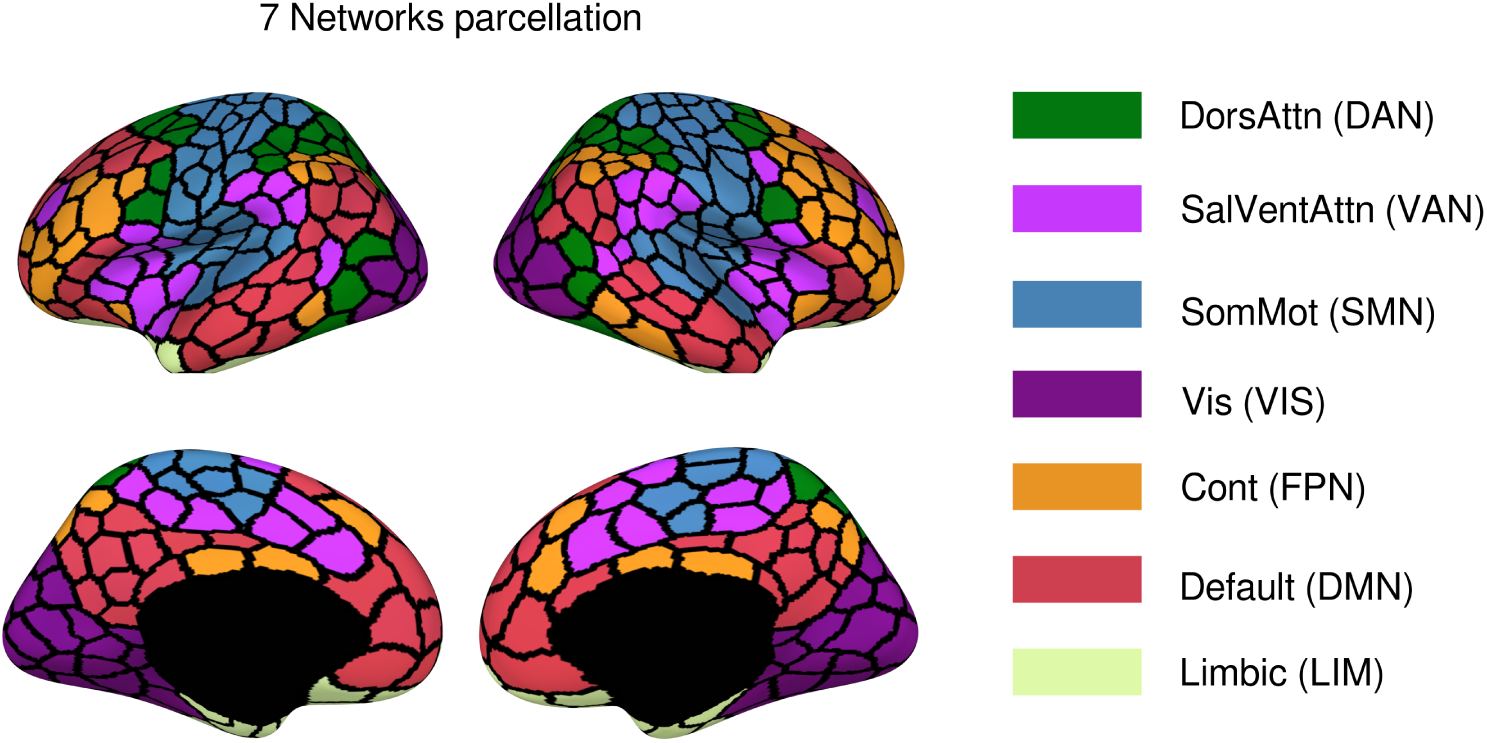
The cortical parcellation (400 parcels) based on the Schaefer atlas.

**Figure S5.**
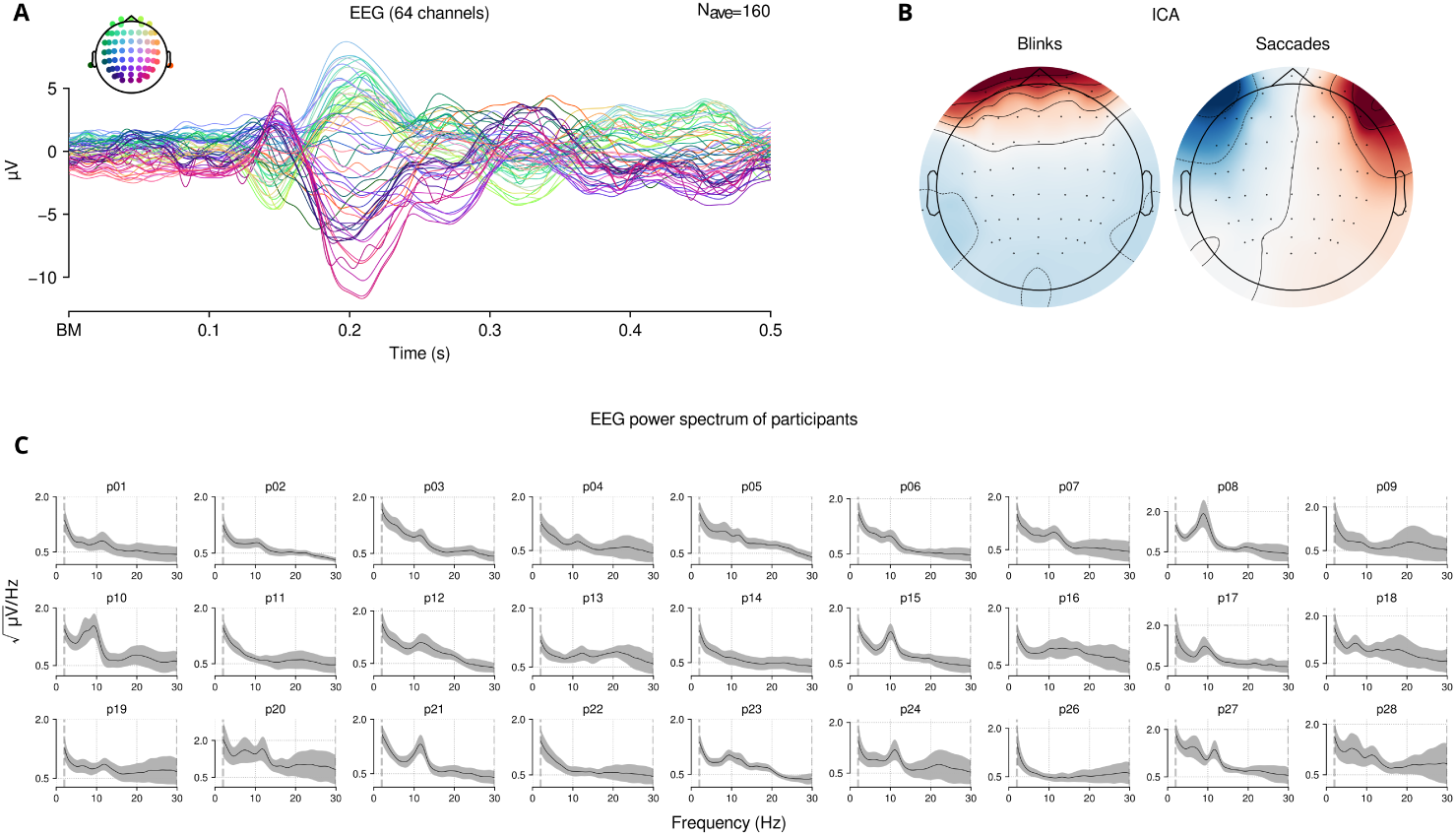
EEG preprocessing steps. **A**) Average evoked activity across electrodes of one participant, from a 500 *ms* time-window starting from the ball movement events. **B**) Template of saccade and eye-blink components which are projected out of the data for each participant, following ICA analysis. **C**) Average power spectra of all participants.

**Figure S6.**
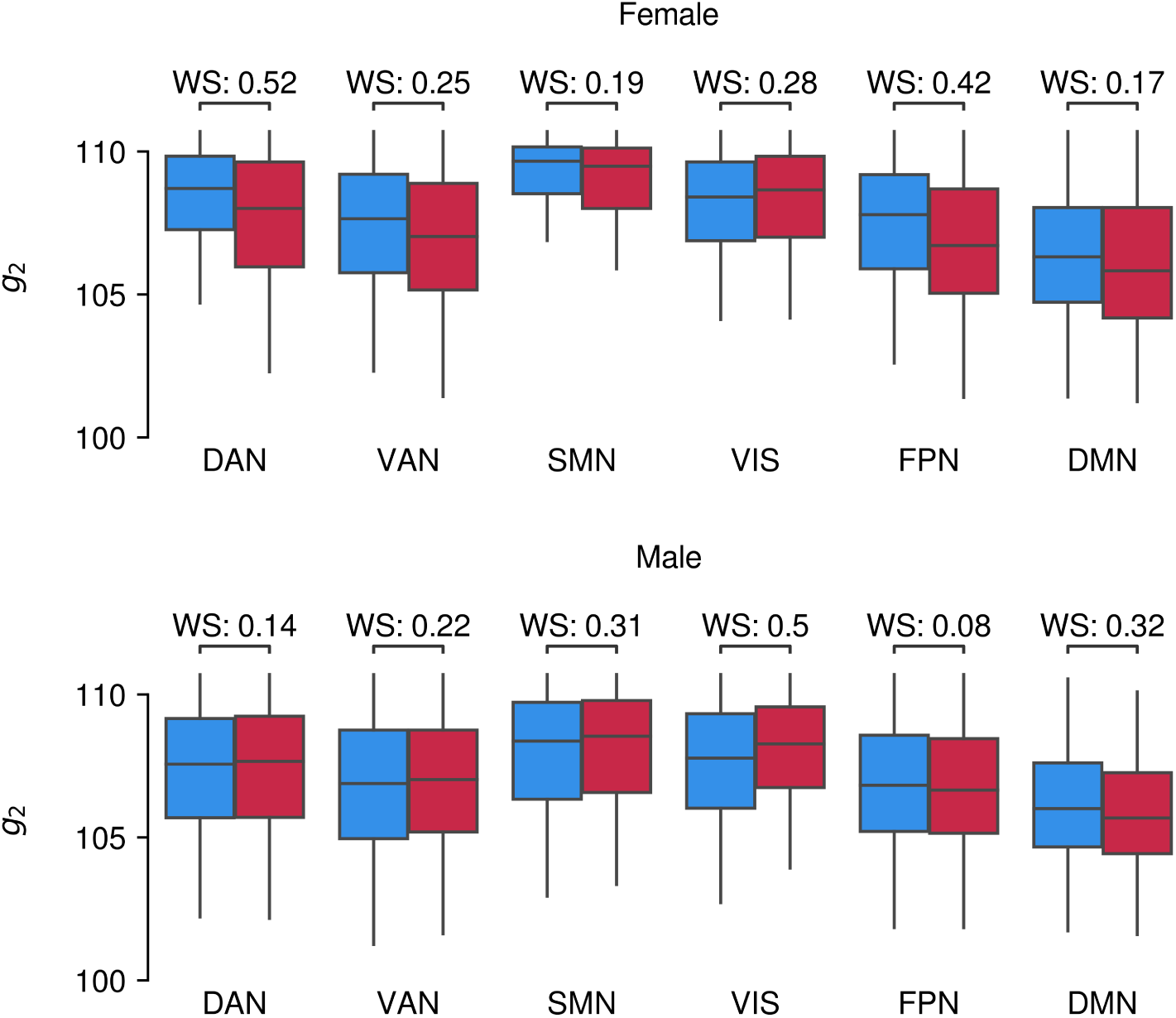
Pooled Distribution of SE at each brain network, across subject groups.

The EEG signals were recorded from all participants throughout the entire experiment. Prior to the start of the experiment, the EEG caps were fitted to each participant’s individual head size. During the process of applying conductive gel to the electrodes and securing the EEG cap, the experimenter began introducing a cover story, mentioning potential slider defects and the possibility of someone entering the room to ‘read the slider resistance’. Before the actual experiment commenced, the experimenter informed the participant that she would leave the room ‘to look for technical support to test the equipment’. Participants were told that the experiment would run entirely without interference from the experimenter, meaning they could continue uninterrupted. Between the 10-trial practice block and the first experimental block, a 30-second break was programmed to allow a time to enter the room (in the case where the first block was the ‘presence’ condition). Similarly, an automatic 2-minute break was inserted between the two experimental blocks, during which the spectator could either enter or leave the room. The presence and absence conditions of the experimental blocks were counterbalanced, meaning that the spectator sometimes entered during the first block and sometimes during the second block. All female participants encountered the same female spectator, while all male participants encountered the same male spectator. Participants were instructed beforehand not to interact with the spectator, as this could introduce noise into the EEG signal.

### 6.5. Inference validation

SBI framework across all scales was validated via in-silico observation, as shown in Supplementary Figure S7, Figure S8, Figure S9). Crucially, due to our amortized approach [28, 29], the trained network can be directly applied to make inference on empirical recordings. This eliminates the need for repeated training for each new data point, leading to significant efficiency gains. The validation has also been applied to non-parametric Hamiltonian Monte Carlo (HMC) sampling, but only at the mesoscopic level (see Supplementary Figure S11), since it becomes computationally prohibitive at the whole-brain scale and impractical at the microscale due to the divergence of such gradient-based methods given the discrete nature of spikes.

**Figure S7.**
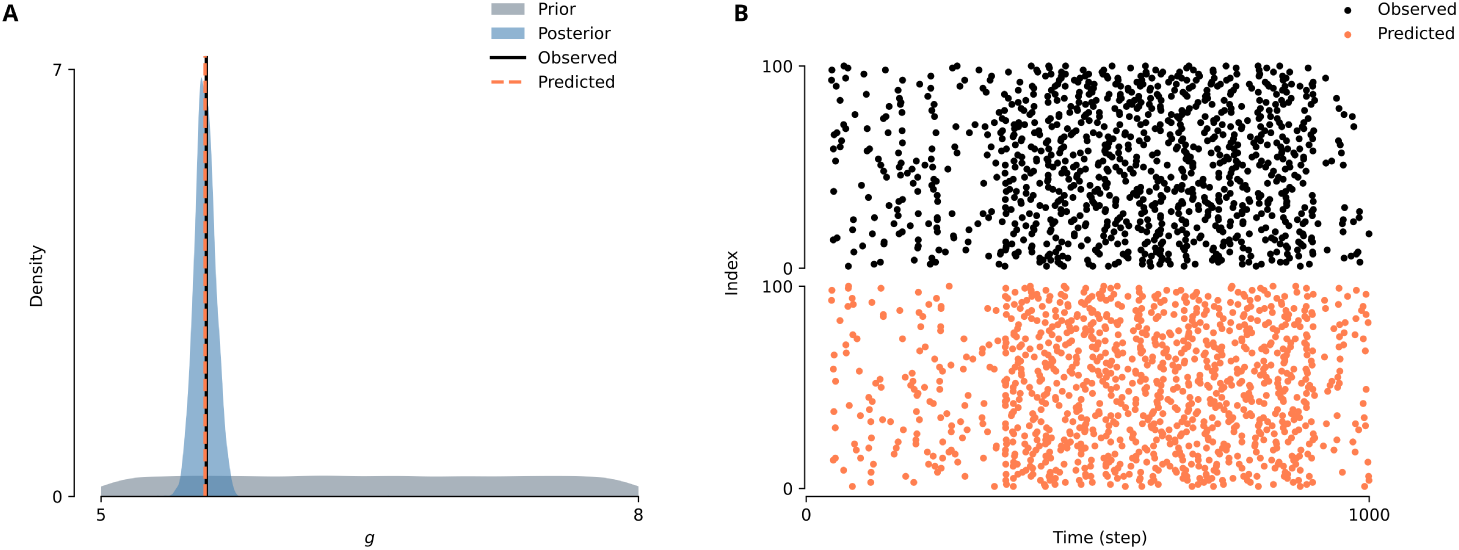
SBI validation at the microscale model. **A**) Prior distribution (in gray) and posterior distribution (in blue) given the ground truth (in black), versus the predicted value of *g* (in orange). **B**) The observed spike rasters generated with ground truth (upper panel) versus predicted (lower panel) generated with posterior samples. Here we used a budget of *n* = 10000 random simulations and maximum firing rate was chosen as the low-dimensional data features for training the neural spline flows.

**Figure S8.**
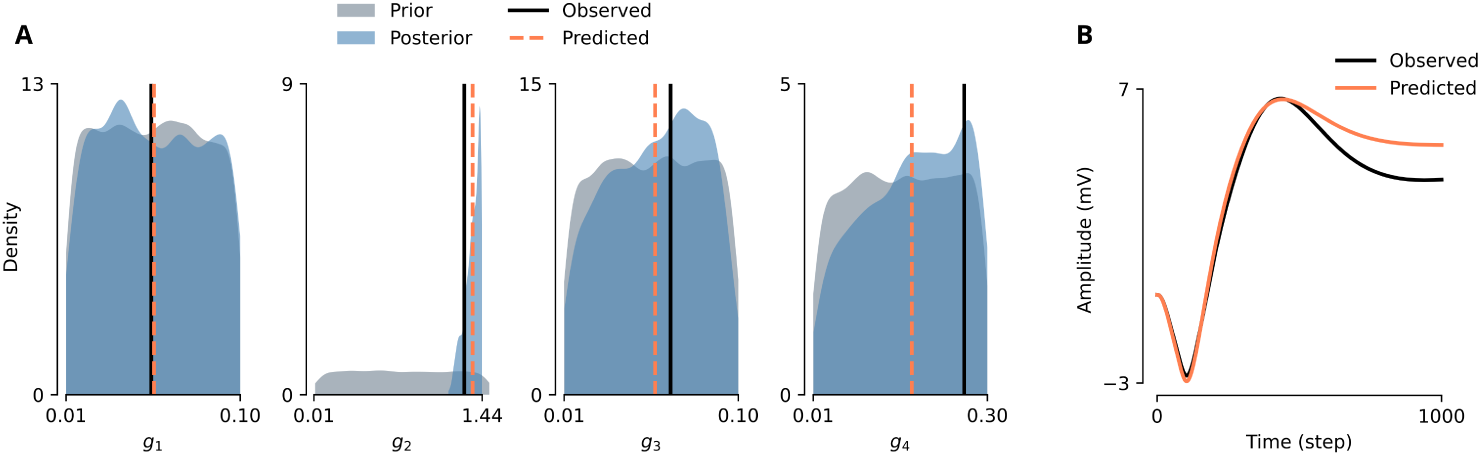
SBI validation at the mesoscale model. **A**) Prior distribution (in gray) and posterior distribution of effective connectivities *g*1−4 (in blue) given the ground truth (in black), versus the predicted parameter values (in orange). We observe a large posterior shrinkage (i.e., Bayesian learning) for *g*2, compared to the other parameters. **B**) The observed ERPs generated with ground truth (in black) versus predicted (in orange) generated with posterior samples. Here we used a budget of *n* = 10000 random simulations, and the peak ERP amplitude values were again selected as the low-dimensional data features for training the neural spline flows.

**Figure S9.**
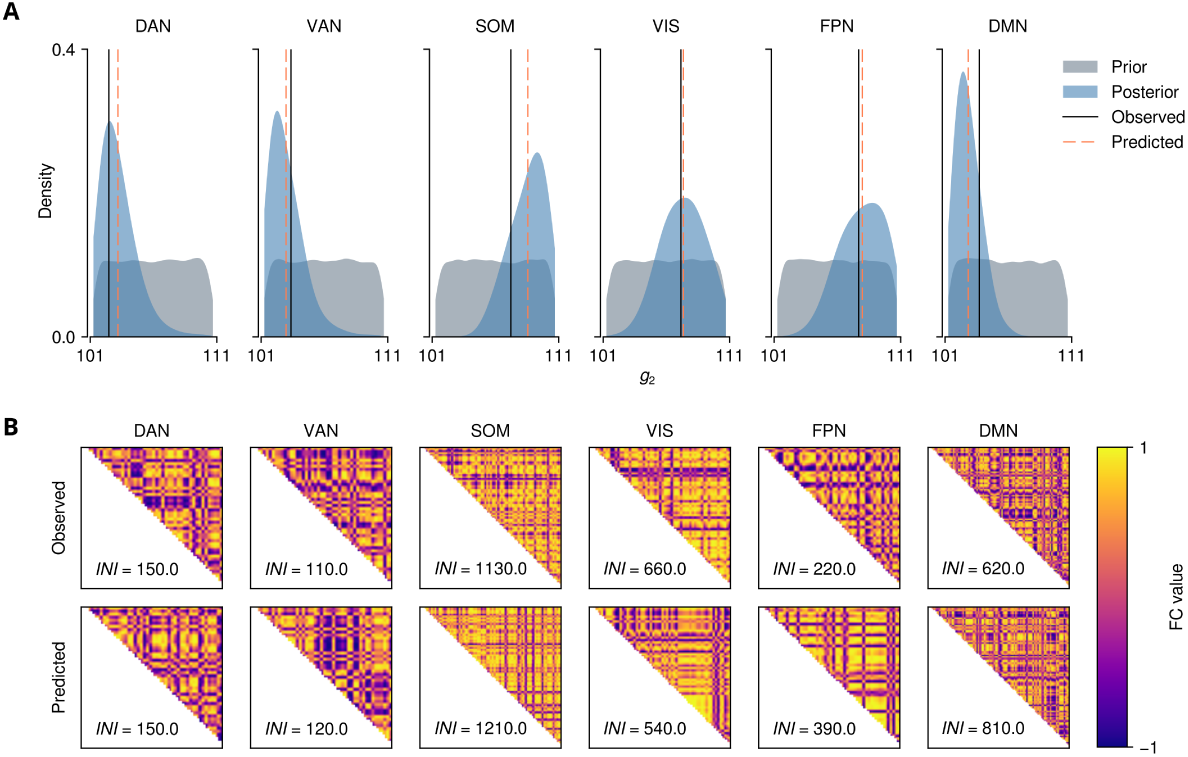
SBI validation at the macroscale model. **A**) Prior distribution (in gray) and posterior distribution of *g*2 (in blue) for the functional networks, given the ground truth (in black), versus the predicted parameter values (in orange). **B**) The observed FC at each network, alongside their respective INI, generated with ground truth (upper panels) versus predicted (lower panels) generated with posteriors samples. Here we used a budget of *n* = 20000 random simulations and INI values were used as the low-dimensional data features for training the neural spline flows.

**Figure S10.**
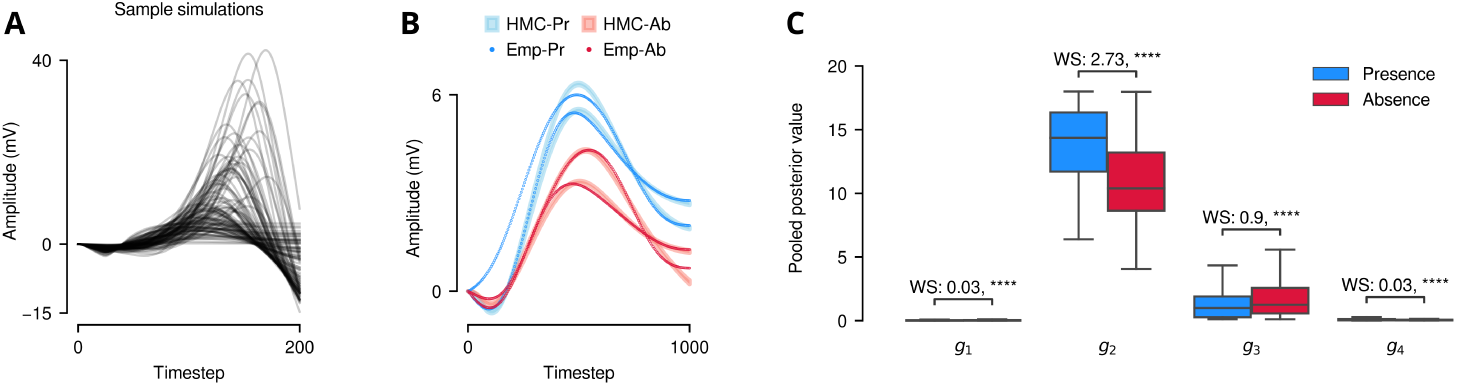
Inference of mesoscale synaptic efficacies using Hamiltonian Monte Carlo (HMC) sampling. **A**) Sample simulations obtained from drawing parameters randomly from the prior distribution. **B**) Fit of observed ERPs from two sessions. **C**) Posterior distribution of SE in the mesoscale model.

**Figure S11.**
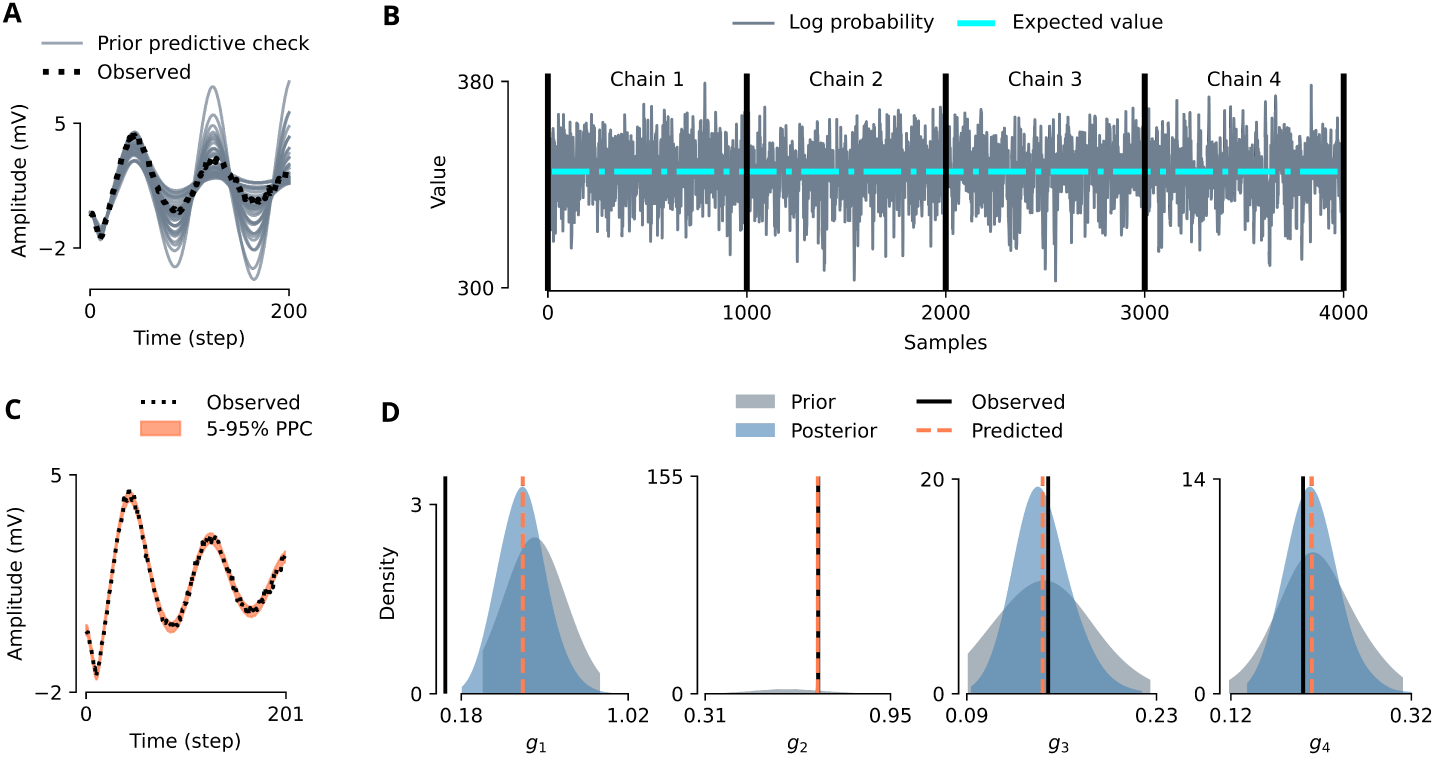
**A**) Prior predictive check (simulations using samples drawn from prior, in grey) versus the observed data (black). **B**) Log probability of the four HMC chains, ran with random initial conditions, converged to the same stationary distribution. **C**) Posterior predictive check (simulations using samples drawn from posterior, in orange) versus the observed data (black). **D**) Estimated posterior distributions using HMC. We observe a large posterior shrinkage (i.e., Bayesian learning) for *g*2, compared to the other parameters.

### 6.6. MCMC-based inference of condition-ERPs

Inferring multiple measures of effective connectivity concurrently presents a significant challenge, as the biological plausibility of the Jansen-Rit model translates to complex relationship between parameter, such as nonlinear degeneracy, and hence multi-modal distributions. Although this type of degeneracy benefits the brain by providing more degrees of freedom, adaptation, and resilience, it poses a waste of computational effort from an inference perspective. However, the use of mean-field approximations in neural mass models (such as sigmoid functions for population firing rates) makes the likelihood function tractable at mesoscale, hence enabling the feasibility of non-parametric inference using Markov chain Monte Carlo (MCMC) sampling. Nevertheless, the efficiency of MCMC sampling methods such as Hamiltonian Monte Carlo (HMC; [105]) is highly sensitive to user-specified algorithm parameters. Therefore, we employed an adaptive version of HMC algorithm known as the No-U-Turn Sampler (NUTS; [106]) to address this challenge. NUTS uses a recursive algorithm to adaptively determine the trajectory of the Markov chain during sampling. This adaptive approach, in conjunction with gradient information from the target posterior density, facilitates efficient exploration of the target distribution in high-dimensional spaces that may exhibit strong correlations [107, 51]. To alleviate the computational cost associated with sampling using NUTS, we leveraged JAX (from NumPyro [108]), a high-performance numerical computation library specializing in composable function transformations and automatic differentiation. With the use of efficient parameterization and weakly informative prior, JAX’s capabilities substantially accelerated our simulations, enabling NUTS sampling of ERPs to converge in less than 1 minute. Using a neural mass model of ERPs (Methods subsection 5.11), effective connectivity parameters were drawn from a gamma distribution Γ(*α, β*) with:

For each ERP, we run 4 chains with *n_warmup_* = 2000 as transitory stage to learn the relation between parameters, and *n_samples_* = 1000 for sampling. The hyper parameters were set as a max tree depth of 12 and target acceptance probability of 0.6. After monitoring the convergence diagnostics (Gelman-Rubin *R*^^^; [109]), the median of the posterior distributions obtained for each ERP was used to calculate the latent space trajectories and manifold embeddings (Methods subsection 5.5). Not only HMC was able to provide excellent fits to our empirical ERPs (Supplementary Figure S10**B**), but the posterior distribution of our chosen SE parameters closely resembled that of SBI (Supplementary Figure S10**C**). The slight difference in *g*_3_ stem from the fact that HMC samples from parameter distributions by going through the timesteps of the time series one by one, without explicit focus on data features relevant to the experimenter (such as maximum amplitude), and therefore obtains parameter distributions that best represent the entirety of the time series.

